# P38 kinases mediate NLRP1 inflammasome activation after ribotoxic stress response and virus infection

**DOI:** 10.1101/2022.01.24.477423

**Authors:** Lea-Marie Jenster, Karl-Elmar Lange, Sabine Normann, Anja vom Hemdt, Jennifer D. Wuerth, Lisa D.J. Schiffelers, Yonas M. Tesfamariam, Florian N. Gohr, Laura Klein, Ines H. Kaltheuner, Dorothee J. Lapp, Jacob Mayer, Jonas Moecking, Hidde L. Ploegh, Eicke Latz, Matthias Geyer, Beate M. Kümmerer, Florian I. Schmidt

## Abstract

Inflammasomes integrate cytosolic evidence of infection or damage to mount inflammatory responses. The inflammasome sensor NLRP1 is expressed in human keratinocytes and coordinates inflammation in the skin. We found that diverse stress signals converge on the activation of p38 kinases to initiate human NLRP1 inflammasome assembly: UV irradiation and microbial molecules that initiate the ribotoxic stress response critically relied on the MAP3 kinase ZAKα to activate p38 and ultimately human NLRP1. Infection with insect-transmitted alphaviruses, including Semliki Forest, Ross River, and Chikungunya virus, also activated NLRP1 in a p38-dependent manner. In the absence on ZAKα, inflammasome assembly was maintained, although at reduced levels, indicating contribution of other upstream kinases. NLRP1 activation by direct nanobody-mediated ubiquitination was independent of p38 activity. Stimulation of p38 by overexpression of MAP2 kinases MKK3 or MKK6 is sufficient for NLRP1 activation, and NLRP1 is directly phosphorylated by p38. Taken together, we define p38 activation as a unifying signaling hub that controls NLRP1 inflammasome activation by integrating a variety of cellular stress signals relevant to the skin.

## Introduction

Inflammatory responses to counteract infection or tissue damage are orchestrated by a multi-layered network of locally and systemically acting signaling cascades. Nucleotide-binding domain and leucine-rich repeat (LRR) containing proteins (NLRs) are critical for the detection and regulation of the early innate immune response in mammalian cells. NLRP1, NLRP3 and other family members initiate the formation of multi-protein-complexes described as inflammasomes (Broz and Dixit, 2016; Mitchell et al., 2019; Tschopp et al., 2003). In response to sensor-specific stimulation, they recruit and nucleate polymerization of the adaptor protein ASC, resulting in micrometer-sized ASC specks within the cell that represent potent caspase-1-activating platforms (Broz et al., 2010). Mature caspase-1 processes the proinflammatory cytokines interleukin (IL)-1β/IL-18 and the pore-forming protein gasdermin D (GSDMD), which leads to pyroptotic cell death (Broz and Dixit, 2016).

Human NLRP1 exhibits a unique domain structure and contains both an N-terminal pyrin domain (PYD) and a C-terminal caspase recruitment domain (CARD) (Martinon et al., 2002; Yu et al., 2017). Autoproteolytic processing of NLRP1 in the ‘function to find domain’ (FIIND) is required for inflammasome formation as it generates the FIIND^UPA^-CARD fragment, which can recruit ASC to initiate the inflammasome response (D’Osualdo et al., 2011; Finger et al., 2012; Gong et al., 2021; Robert Hollingsworth et al., 2021; Sandstrom et al., 2019). In steady-state, FIIND^UPA^-CARD remains associated with the N-terminus; FIIND^UPA^-CARD released from the N-terminus can also be sequestered by association with DPP9 (Hollingsworth et al., 2021; Huang et al., 2021; Okondo et al., 2018; Zhong et al., 2018). FIIND^UPA^-CARD is released by the inhibition of DPP9 (Okondo et al., 2018; Zhong et al., 2018) or by degradation of the NLRP1 N-terminus. The latter activation mechanism enables NLRP1 to detect pathogenic protease activity, as shown for enteroviral proteases (Robinson et al., 2020; Tsu et al., 2021). Other reported stimuli of NLRP1 are ATP depletion (Liao and Mogridge, 2013), UV irradiation (Feldmeyer et al., 2007; Fenini et al., 2018a), Semliki Forest virus (SFV) infection, and dsRNA (Bauernfried et al., 2020). How signaling by these diverse triggers converge, and how they activate NLRP1, is only partly understood.

In rodent Nlrp1b, the N-terminal PYD is replaced by unrelated sequences (Yu et al., 2017). While activation by Dpp9 inhibition is shared with human NLRP1 (Gai et al., 2019; de Vasconcelos et al., 2019), Nlrp1b can be activated by unique triggers such as anthrax toxin lethal factor (Levinsohn et al., 2012), but does not respond to SFV or dsRNA (Bauernfried et al., 2020). The human inflammasome sensor CARD8 exhibits an autoproteolytically processed FIIND^UPA^-CARD fragment similar to NLRP1 and represents the dominant sensor of DPP9 inhibition in a variety of cell types and cell lines (Johnson et al., 2018). Importantly, CARD8 FIIND^UPA^-CARD directly engages pro-caspase-1 and is not able to assemble ASC specks (Gong et al., 2021).

In human keratinocytes, NLRP1, but not CARD8, assembles functional inflammasomes (Bauernfried et al., 2020; Burian and Yazdi, 2018; Fenini et al., 2018a). The prominent role of NLRP1 in the skin is underlined by strong pathological manifestations in the skin observed in patients bearing gain-of-function mutants of NLRP1 (Herlin et al., 2019; Zhong et al., 2016). In this study, we analyzed the signaling cascades upstream of NLRP1 and investigated whether different signaling inputs employ a common mechanism to initiate NLRP1 activation.

## Results

### Robust quantification of human NLRP1 inflammasome assembly by flow cytometry

To identify and evaluate (patho)physiological triggers of NLRP1, we used two complementary cellular systems: In a bottom-up approach, we reconstituted NLRP1 inflammasome components in human embryonic kidney (HEK) 293T cells (Fig. 1). To study human NLRP1 at endogenous levels in a physiologically relevant cell type, we used immortalized N/TERT-1 keratinocytes (Fig. 2). As a readout for activation, we decided to evaluate the assembly of ASC specks, as all downstream effects of NLRP1 rely on the formation of these large macromolecular assemblies, and as CARD8 is not able to initiate ASC specks. Formation of ASC specks is the most proximal event to NLRP1 activation that can be robustly detected and, importantly, is not regulated at the transcriptional level and thus insensitive to most indirect effects.

**Fig. 1.**
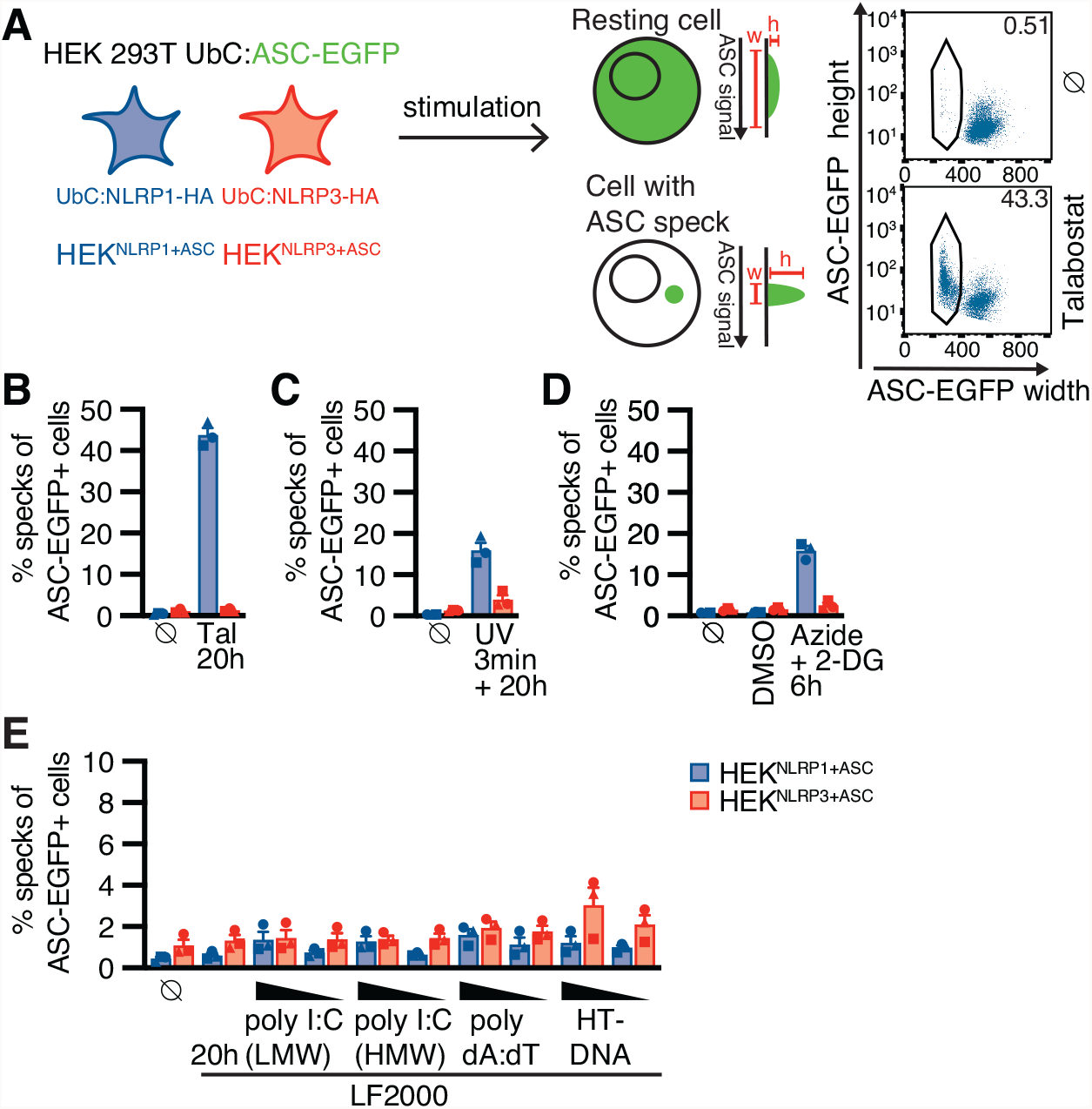
Reconstituted reporter cell lines recapitulate NLRP1 inflammasome assembly. (**A**) Scheme of generated reporter cell lines and detection of ASC specks by flow cytometry. (**B-E**) HEK^NLRP1+ASC^ or HEK^NLRP3+ASC^ cells were treated with the indicated stimuli for 20 h (B-C, E) or 6 h (D). ASC-EGFP-positive cells were analyzed by flow cytometry and the fraction of cells with ASC specks was determined with the gating strategy described in (A). Cells were treated with 30 µM talabostat (Tal) (B), UV for 3 min (C), 10 mM sodium azide and 50 mM 2-deoxyglucose (2-DG) (D), or transfected with 1 or 0.2 µg/mL of the indicated nucleic acid species (E). Data represents average values (with individual data points) from three independent experiments ± SEM; UbC indicates ubiquitin C (promoter), LF2000 indicates Lipofectamine 2000, HT-DNA indicates herring testis DNA.

**Fig. 2.**
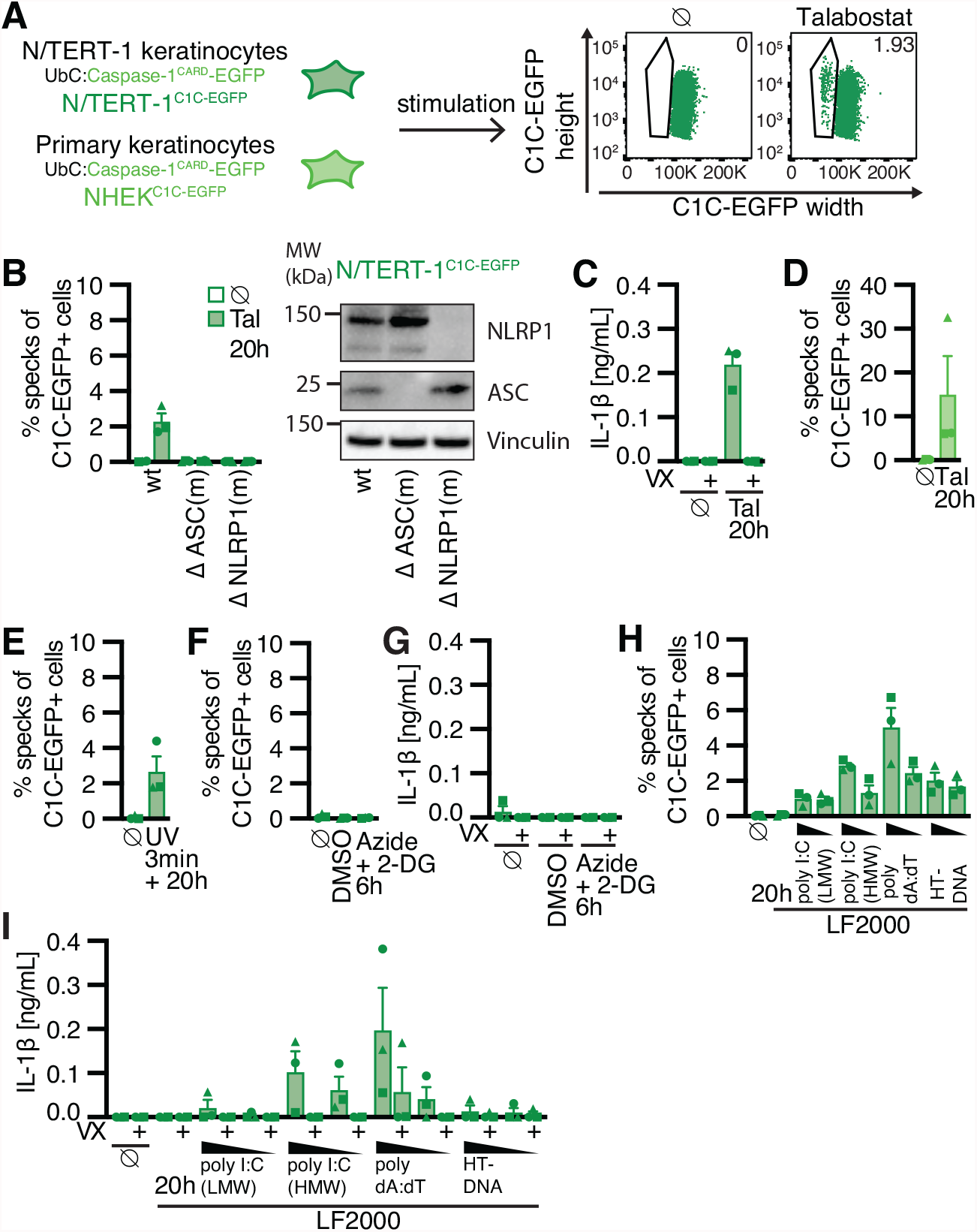
Reporter keratinocyte cell lines recapitulate NLRP1 inflammasome assembly. (**A**) Scheme of generated N/TERT-1 reporter cell lines and detection of caspase-1^CARD^ (C1C) specks by flow cytometry. N/TERT-1^C1C-EGFP^ cells and their derivatives (B, C, E-I) or primary normal human epidermal keratinocytes expressing C1C-EGFP (NHEK^C1C-EGFP^) (D) were treated with the indicated stimuli for 20 h (B-E, H, I) or 6 h (F, G). To quantify inflammasome assembly, N/TERT-1 derivates were stimulated in the presence of 100 µM Vx-765 (VX) and C1C-EGFP-positive cells were analyzed by flow cytometry. The fraction of cells with C1C-EGFP specks was determined with the gating strategy described in A (B, D-F, H, I). IL-1β from the supernatants of cells stimulated in the absence or presence of VX was quantified by Homogeneous Time Resolved Fluorescence (HTRF) (C, G, I). (**B, C**) Wildtype N/TERT-1^C1C-EGFP^ and the indicated monoclonal knockouts of ASC or NLRP1 were treated with 30 µM talabostat and analyzed for C1C-EGFP specks (B) and IL-1β secretion (C); knockouts were confirmed by immunoblot with the indicated antibodies (B). (**D**) Primary NHEK^C1C-EGFP^ were treated with 30 µM talabostat and analyzed for C1C-EGFP specks. N/TERT-1^C1C-EGFP^ were treated with UV for 3 min (**E**), 10 mM sodium azide and 50 mM 2-DG (**F, G**), or transfected with 1 or 0.2 µg/mL of the indicated nucleic acid species (**H, I**), and analyzed for C1C specks (E, F, H) or IL-1 β secretion (G, I). Data represents average values (with individual data points) from three independent experiments ± SEM.

We equipped a monoclonal HEK 293T cell line expressing ASC-EGFP from a weak ubiquitin promoter (pUbC) with either NLRP1 or NLRP3, which was likewise under the control of pUbC (Fig. 1A). We next selected clones with optimal signal to noise ratio (HEK^NLRP1+ASC^ and HEK^NLRP3+ASC^), and quantified assembly of ASC specks by flow cytometry, exploiting the characteristic redistribution of ASC-EGFP into a single speck per cell: Cells with ASC-EGFP specks passing the detectors of the flow cytometer cause a characteristic EGFP signal with decreased width and increased height, compared to untreated cells with diffuse ASC-EGFP, which show a broader, less intense EGFP signal (Sester et al., 2015) (Fig. 1A). To validate the cell lines, we treated them with the DPP9 inhibitor talabostat (Tal) (Fig. 1, A and B, and fig. S1A). More than 40% of the cells expressing NLRP1 assembled ASC specks in response to talabostat, while none of the NLRP3-expressing cells showed a response. HEK^NLRP3+ASC^ assembled ASC specks after treatment with LPS and nigericin, validating a functional NLRP3 inflammasome (fig. S1B).

We next tested whether UV treatment as a common environmental threat to keratinocytes activated NLRP1 as postulated before (Fenini et al., 2018a). We found that UV treatment of HEK^NLRP1+ASC^ cells, but not HEK^NLRP3+ASC^ cells, indeed induced the assembly of inflammasomes, which were robustly detectable after 20 h (Fig. 1C). Depletion of cellular energy levels had likewise been proposed to activate NLRP1 (Liao and Mogridge, 2013). ATP depletion in reporter cells with azide and 2-deoxyglucose (2-DG) rapidly led to NLRP1-dependent ASC specks (Fig. 1D). Due to the strong toxicity of the treatment to the cells, inflammasome assembly was quantified after 6 h. Cytosolic double-stranded RNA (dsRNA) was recently proposed to activate NLRP1 as a direct ligand (Bauernfried et al., 2020). We therefore transfected reporter cells with the synthetic dsRNA poly(I:C), or double-stranded DNA in the form of synthetic poly(dA:dT) or herring testis DNA (HT-DNA). Cytosolic delivery of neither molecule was able to robustly activate NLRP1 or NLRP3 inflammasomes above background levels (Fig. 1E).

To assess human NLRP1 inflammasome assembly in keratinocytes expressing endogenous levels of NLRP1 and ASC, we equipped N/TERT-1 keratinocytes known to express NLRP1 (Zhong et al., 2016) with a reporter construct composed of the caspase recruitment domain (CARD) of caspase-1 (C1C) fused to EGFP. C1C-EGFP is efficiently recruited to ASC specks, allowing visualization and quantification of inflammasome assembly by fluorescence microscopy and flow cytometry, exploiting the characteristic redistribution of fluorescence into a single speck per cell as for ASC-EGFP (Fig. 2A, and fig. S2A). Unlike fluorescent fusions of ASC, however, this reporter does not alter the endogenous levels of ASC, cannot assemble specks by mere overexpression, and lastly also recapitulates the recruitment of caspase-1. All experiments for the quantification of specks were performed in the presence of the caspase-1 inhibitor Vx-765 (VX) to avoid loss of responding cells by pyroptosis. The novel reporter will be described and characterized in greater detail in an independent study (in preparation). Importantly, N/TERT-1^C1C- EGFP^ cells exhibited robust inflammasome assembly after talabostat stimulation and no specks were observed in cells lacking ASC or NLRP1 (Fig. 2, A and B). Only around 2% of keratinocytes assembled ASC specks in response to talabostat, which may be attributed to the endogenous protein levels or a higher threshold of activation. Importantly, no specks whatsoever were observed in the absence of activation, which allowed us to confidently quantify inflammasome triggers with low response rates. N/TERT-1 keratinocytes express pro-IL-1β and can therefore secrete caspase-1-matured IL-1β upon inflammasome activation. We found that secretion of mature IL-1β correlated well with the observed speck response (Fig. 2C). Primary Normal Human Epidermal Keratinocytes (NHEK) can be efficiently transduced with lentiviruses. This allows us to equip those cells with C1C-EGFP as well to validate the most important findings in primary cells. NHEK robustly assembled ASC specks in response to talabostat treatment (Fig. 2D). We next tested other described NLRP1 activators and found that N/TERT-1^C1C-EGFP^ cells assembled inflammasomes upon UV treatment and exhibited a response rate comparable to talabostat (Fig. 2E). No ASC specks or IL-1β secretion were detected in response to azide and 2-DG, although it cannot be ruled out that this is caused by the rapid cell death upon ATP depletion (Fig. 2, F and G). Unlike HEK^NLRP1+ASC^, N/TERT-1^C1C-EGFP^ assembled inflammasomes and secreted IL-1β when transfected with dsRNA as well as DNA (Fig. 2, H and I), suggesting that additional factors absent in HEK 293T cells contributed to NLRP1 activation.

The established series of reporter cells robustly recapitulated NLRP1 inflammasome assembly combined with simple quantification by flow cytometry. HEK-based reporter cells that only differ by the expressed inflammasome sensor NLRP1 or NLRP3 can help to distinguish whether any treatment affects inflammasome assembly in general, or NLRP1 inflammasomes specifically. The HEK reporters exhibit a high fraction of responding cells and we found that they are less sensitive to treatments with potentially cytotoxic effects. N/TERT-1-based reporter cells reflect endogenous protein levels in a cell type relevant for NLRP1 inflammasomes and recapitulate the entire inflammatory signaling pathway.

### Direct targeting of NLRP1 PYD for ubiquitination is sufficient for activation

Murine Nlrp1b (mNlrp1b) is activated by proteolytic cleavage of the N-terminus by *B. anthracis* lethal factor (Chavarria-Smith and Vance, 2013; Levinsohn et al., 2012), while human NLRP1 is cleaved and activated by viral proteases (Robinson et al., 2020; Tsu et al., 2021). This is followed by proteasomal degradation of the destabilized N-terminus, releasing the C-terminal FIIND^UPA^-CARD fragment sufficient for inflammasome assembly (Chui et al., 2019; Sandstrom et al., 2019). mNlrp1b was also shown to be directly activated by a ubiquitin ligase of the human pathogen *Shigella flexneri* (Sandstrom et al., 2019). To test if ubiquitination of human NLRP1 with its distinct N-terminus was also sufficient for activation, we sought to engineer an experimental setup to specifically ubiquitinate human NLRP1 at will. We immunized alpacas with the PYD of NLRP1 and employed phage display to identify NLRP1^PYD^-specific variable domains of heavy chain-only antibodies (VHH), also described as nanobodies (fig. S3A). Specific binding of VHH_NLRP1 PYD_ 1 (VHH_PYD_ 1) and VHH_NLRP1 PYD_ 2 (VHH_PYD_ 2) to NLRP1^PYD^ was confirmed by ELISA (fig. S3B). Unlike antibodies, nanobodies often function in the reducing environment of the cytosol. They can thus be genetically fused to enzymes to recruit their activity to the target proteins of nanobodies. To explore whether NLRP1^PYD^-specific nanobodies allow activation of NLRP1 by targeted ubiquitination, we next generated expression vectors for fusions of nanobodies to the human ubiquitin ligase receptor von Hippel Lindau (VHL) (Fig. 3A). Endogenous VHL serves as a ubiquitin ligase receptor for modular Cullin-2 ubiquitin ligases and mediates the constitutive ubiquitination and proteasomal degradation of HIF1α under normoxic conditions (Haase, 2009), suggesting that overexpression as such will not alter cellular states at normal oxygen levels. Overexpressed VHL-VHH fusions have been successfully used to mediate proteasomal degradation of VHH targets (Fulcher et al., 2017).

**Fig. 3.**
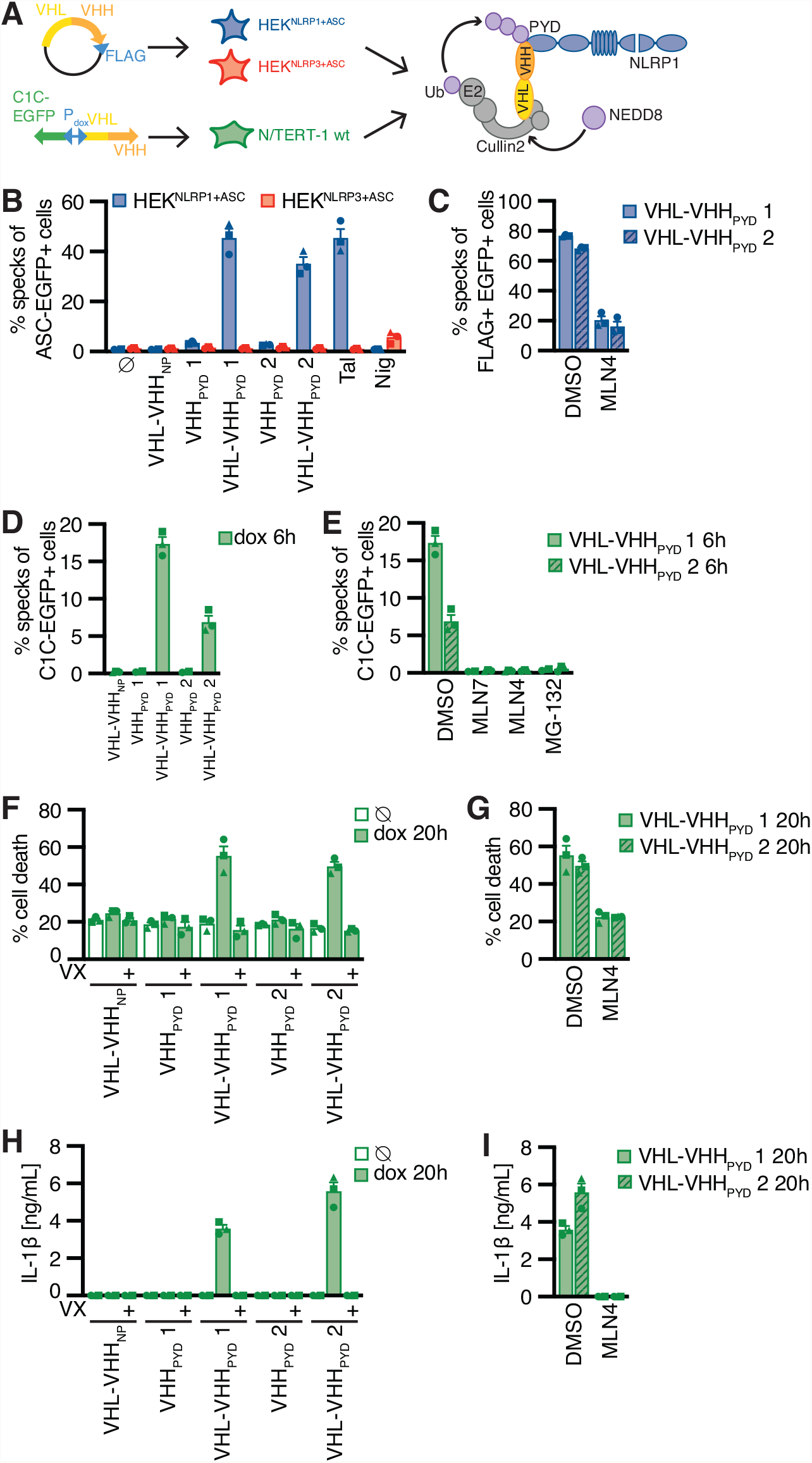
Ubiquitination of NLRP1^PYD^ is sufficient for activation of human NLRP1. **A**) Experimental setup and scheme of nanobody-mediated ubiquitination of NLRP1. HEK^NLRP1+ASC^ and HEK^NLRP3+ASC^ cells were transiently transfected with expression vectors for HA-tagged VHHs or FLAG-tagged VHL-VHH fusions; N/TERT-1 cells were transduced with lentiviral vectors encoding C1C-EGFP and (VHL-)VHH controlled by a bidirectional doxycycline (dox)-inducible promoter. (**B**) HEK^NLRP1+ASC^ and HEK^NLRP3+ASC^ expressing the indicated VHL-VHH fusions or VHHs alone were analyzed for ASC-EGFP specks 20 h post transfection. Control samples were treated with 30 µM talabostat for 20 h or 10 µM Nigericin (Nig) for 1 h. (**C**) HEK^NLRP1+ASC^ were transfected as in (B), treated with 1 µM MLN4924 (MLN4) or DMSO 4 h post transfection, stained for FLAG and analyzed for speck formation in FLAG-positive cells after 20 h. (**D-I**) N/TERT-1 cells inducibly expressing VHL-VHH or VHH alone were treated with 1 µg/mL dox for the indicated time in presence of 100 µM VX (D,E), or in the presence of VX as indicated (F, H), and subsequently analyzed for C1C-EGFP specks (D, E), cell death by lactate dehydrogenase (LDH) release (F, G), or IL-β release by HTRF (H, I). Where indicated, cells were treated with 1 µM MLN4, 1 µM MLN7243 (MLN7), 1 µM MG-132, or DMSO. Data represents average values (with individual data points) from three independent experiments ± SEM.

We next transfected HEK^NLRP1+ASC^ cells with expression vectors for epitope-tagged VHHs or VHL-VHH fusions. Transient overexpression of FLAG-tagged VHL-VHH_PYD_ 1 and VHL-VHH_PYD_ 2 induced ASC speck formation in more than 40% of ASC-EGFP positve cells (Fig. 3B, fig. S3C). When we gated for FLAG-positive cells, almost 80% of the cells exhibited ASC specks (Fig. 3C), indicating near complete activation of NLRP1 in cells expressing the fusion protein, likely independent of the bottlenecks of endogenous NLRP1 activation yielding lower response rates. Importantly, NLRP1 was not activated by overexpression of the nanobodies alone, or by VHL fusions to a control nanobody targeting influenza A virus NP (Ashour et al., 2015) (Fig. 3B). VHL-VHH_PYD_ 1 and VHL-VHH_PYD_ 2 did not activate NLRP3 inflammasomes either. Cullin-based ubiquitin ligase complexes are critically dependent on modification with the ubiquitin-like molecule NEDD8 (Soucy et al., 2009). Inhibition of NEDD8-activating enzyme with MLN4924 (MLN4) 4 h post transfection blocked NLRP1 inflammasome activation without changing the expression levels of VHL-VHH_PYD_ 1 and VHL-VHH_PYD_ 2 (Fig. 3C, fig S3D), confirming that inflammasome assembly in this setup depended on cullin ubiquitin ligase activity. Importantly, unlike the proteasome inhibitors MG-132 and bortezomib as well as the E1 inhibitor MLN7924 (MLN7), MLN4 was not toxic during prolonged treatment up to 20 h.

We next generated N/TERT-1 keratinocyte cell lines expressing C1C-EGFP as well as VHHs or VHL-VHH fusions under the control of a bidirectional doxycycline-inducible promoter. Induced expression of VHL-VHH_PYD_ 1 and VHL-VHH_PYD_ 2 triggered inflammasome assembly detectable as early as 6 h post induction, with faster activation by VHH_PYD_ 1 than by VHH_PYD_ 2 (Fig. 3D, fig. S3E). Again, response rates were substantially higher than after talabostat treatment. Neither nanobody expression alone, nor VHL fusions to the control nanobody VHH_NP-1_ induced inflammasome assembly. The rapid inflammasome assembly allowed us to quantify responses in the presence of inhibitors of E1 ubiquitin activating enzyme (MLN7), the proteasome (MG-132), as well as neddylation (MLN4) (Fig. 3E). We found that inflammasome assembly was completely blocked by either inhibitor, indicating that VHL fusions of NLRP1^PYD^ nanobodies indeed rely on the activation of ubiquitin, cullin ubiquitin ligases, as well the proteasome to mediate ubiquitination and N-terminal degradation of NLRP1. None of the inhibitors affected expression levels of C1C-EGFP (fig. S3F). Assembly of ASC specks was accompanied by both pyroptotic cell death, as assessed by release of lactate dehydrogenase (LDH) (Fig. 3, F and G), and robust IL-1β secretion (Fig. 3, H and I). This confirmed the use of ASC specks as a robust readout for inflammasome responses. Pyroptosis and cytokine secretion were completely abrogated by caspase-1 inhibition with Vx-765 as expected. Likewise, interference with cullin neddylation abrogated all inflammasome responses (Fig. 3, E, G and I). All generated cell lines exhibited comparable levels of pyroptosis and IL-1β release when treated with talabostat in the absence of transgene expression (fig. S3, G and H).

We thus demonstrate that ubiquitination of human NLRP1 is sufficient for its activation by N-terminal degradation. Importantly, this well-controlled system allows the precise direct activation of NLRP1 in the absence of any upstream signal amplification. It neither relies on small molecules with difficult to evaluate specificity, nor on pathogen infection or pathogen-associated molecular patterns that may trigger multiple pathways at the same time. In our efforts to define the signaling cascades of physiological NLRP1 activation, this system will therefore serve as a valuable control to distinguish perturbations that affect NLRP1 inflammasomes directly from those that interfere with the upstream signals initiated by one or several triggers.

### UV treatment activates NLRP1 inflammasomes through p38 kinase activity

We next followed up on NLRP1 activation by UV, as this is a relevant (and known) environmental trigger of inflammation. Human keratinocytes are regularly exposed to UVB, and UV-triggered activation of NLRP1 was well recapitulated in HEK 293T and N/TERT-1 keratinocyte-based reporter cells (Fig. 1C and 2E). UV irradiation damages multiple molecules in the cell directly or indirectly. This includes 1) DNA damage and activation of distinct signaling pathways including kinases ATM and ATR, 2) RNA damage inducing the ribotoxic stress response, as well as 3) the generation of reactive oxygen species (ROS) modifying multiple molecule classes. Induction of DNA damage in HEK^NLRP1+ASC^ reporter cells with doxorubicin and etoposide did not initiate inflammasome assembly beyond background levels, although phosphorylation of γH2AX as readout for ATM activation could be verified by flow cytometry (fig. S4A). Likewise, inhibition of ATM or ATR did not impair UV-induced NLRP1 activation (fig. S4B). While H_2_O_2_ treatment caused NLRP1 inflammasome activation in HEK^NLRP1+ASC^ cells, and even some NLRP3-dependent specks in HEK^NLRP3+ASC^ (fig. S4C), no H_2_O_2_-mediated inflammasome activation or IL-1β secretion was observed in N/TERT-1^C1C-EGFP^ (fig. S4, D and E). This suggested that ROS were not sufficient for the NLRP1 inflammasome activation observed in both cell types.

UV treatment activates the ribotoxic stress response by damaging cellular RNA. This includes damage of ribosomal RNAs that results in clashing ribosomes. This response initiates MAP kinase signaling by activating the mitogen-activated protein kinase kinase kinase (MAP3K) ZAKα (MAP3K20)(Vind et al., 2020a; Wang et al., 2005), which phosphorylates mitogen-activated protein kinase kinases (MAP2K) MKK3 (MAP2K3) as well as MKK6 (MAP2K6) (Vind et al., 2020b). These, in turn, activate the four p38 isoforms p38α (MAPK14), p38β (MAPK11), p38γ (MAPK12), and p38δ (MAPK13) by phosphorylation (Canovas and Nebreda, 2021). p38α is abundantly expressed in many cell types, while p38β is ubiquitously expressed at low levels. P38γ and p38δ exhibit more tissue-restricted expression patterns. To test whether UV-induced inflammasome assembly relied on the ribotoxic stress response and p38 signaling, we quantified speck formation in HEK^NLRP1+ASC^ and N/TERT-1^C1C-EGFP^ cells treated with the p38α/β inhibitor SB202190 (SB), or the pan-p38 inhibitor doramapimod (Dora) (Fig. 4, A and B). Assembly of ASC specks was completely abrogated by both inhibitors, suggesting that UV-induced NLRP1 activation was indeed critically dependent on p38 kinase activity. In line with that, we observed robust phosphorylation of p38 after UV irradiation (Fig. 4C). Inflammasome assembly was not sensitive to ISRIB, an inhibitor of the integrated stress response, and only partially affected by Jnk-In-8 (Jnk), a reversible inhibitor of JNK1, JNK2, and JNK3. When we used PF3644022 (PF) to inhibit MAPKAPK2 (MK2), an effector kinase directly phosphorylated by p38 regulating stability of mRNA transcripts, no changes in NLRP1 activation were observed. This indicates that inflammasome activation was not caused by altered mRNA stability, although p38-dependent gene regulation could not be ruled out.

**Fig. 4.**
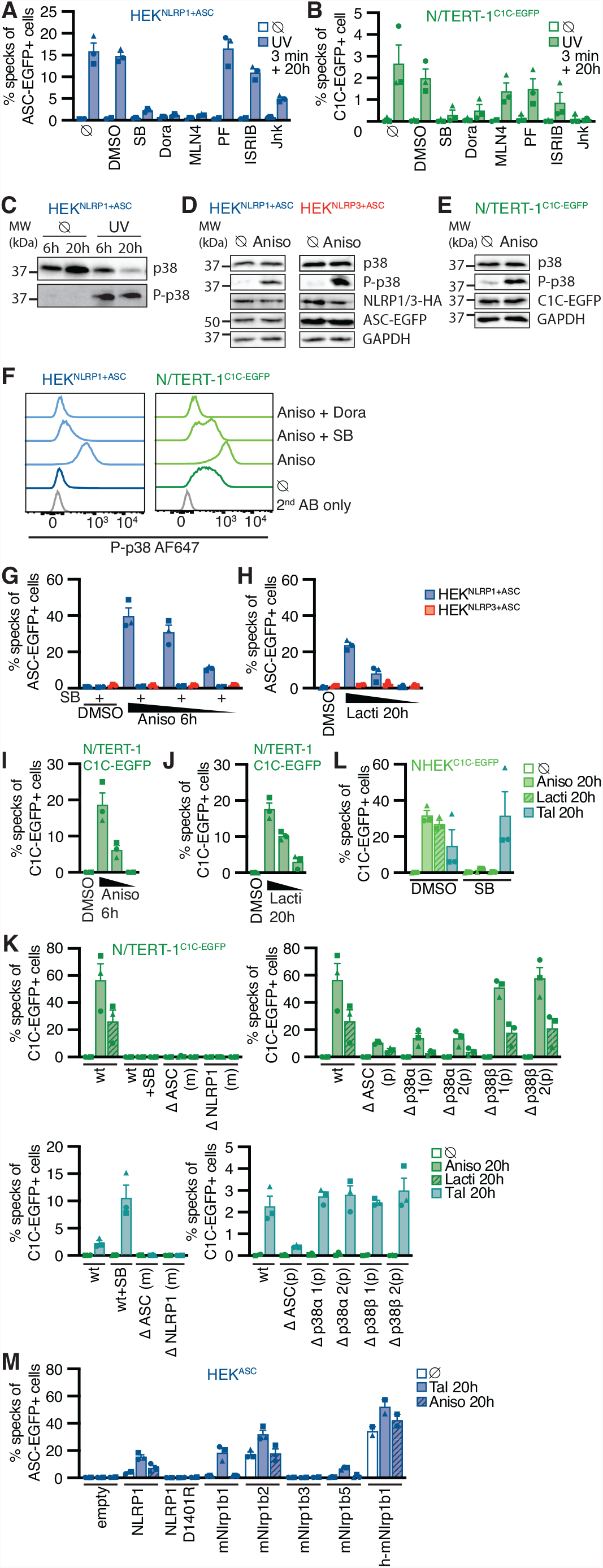
Human NLRP1 is activated by the ribotoxic stress response. (**A-C**) HEK^NLRP1+ASC^ (A, C) or N/TERT-1^C1C-EGFP^ (B) cells were treated with UV for 3 min and cultivated for 20 h in the presence of 20 µM SB202190 (SB), 10 µM Doramapimod (Dora), 1 µM MLN4, 1 µM PF3644022 (PF), 200 nM ISRIB, 3 µM Jnk-In-8 (Jnk), or DMSO. Speck formation was quantified by flow cytometry (A, B) or lysates analyzed by immunoblot with antibodies for p38 and phospho-p38 (P-p38) (C). **(D-F)** HEK^NLRP1+ASC^ (D, F), HEK^NLRP3+ASC^ (D), or N/TERT-1^C1C-EGFP^ (E, F) were treated with 15 µM anisomycin (Aniso) for 60 min. Lysates were analyzed by immunoblot with antibodies for p38, P-p38, HA, EGFP, or GAPDH (D, E). Fixed cells were stained for P-p38 and analyzed by flow cytometry (F). (**G, I**) HEK^NLRP1+ASC^, HEK^NLRP3+ASC^ (G) or N/TERT-1^C1C-EGFP^ (I) cells were treated with 15, 1.5, or 0.15 µM anisomycin for 6 h, where indicated in the presence of 20 µM SB. Specks were quantified as in (A). **(H, J)** HEK^NLRP1+ASC^ or HEK^NLRP3+ASC^ (F) or N/TERT-1^C1C-EGFP^ (J) cells were treated with 2.5, 0.5, and 0.1 µM lactimidomycin (Lacti) for 20 h and analyzed as in (A). (**K**) N/TERT-1^C1C-EGFP^ wildtype cells, or the indicated monoclonal (m) or polyclonal (p) knockout cell lines were treated with 15 µM anisomycin for 6 h or 30 µM talabostat for 20 h, where indicated in the presence of 20 µM SB. Cells were analyzed as in (A). (**L**) NHEK^C1C-EGFP^ were stimulated with 15 µM anisomycin, 2 µM lactimidomycin, or 30 µM talabostat in the presence of DMSO or 20 µM SB, and analyzed as in (A). (**M**) HEK^ASC^ cells transiently expressing human NLRP1, the indicated murine Nlrp1b alleles, or a chimeric protein comprised of PYD and linker of NLRP1 (aa 1-327) and NACHT-LRR-FIIND-CARD of mNlrp1b (aa 126-1233), were treated with 30 µM talabostat or 15 µM anisomycin. Cells were analyzed as in D. Data from all flow cytometry experiments quantifying specks represents average values (with individual data points) from three independent experiments ± SEM. Immunoblots in (C-E), as well as flow cytometry data in (F) display experiments representative of three independent experiments. N/TERT-1^C1C-EGFP^ and NHEK^C1C-EGFP^ cells were stimulated in the presence of 100 µM VX for all flow cytometry experiments.

### The ribotoxic stress response triggers NLRP1 activation

We next evaluated whether other activators of the ribotoxic stress response were also able to activate NLRP1 inflammasomes. We first tested anisomycin (Aniso), an antibiotic produced by *Streptomyces griseolus*, which inhibits protein synthesis by interfering with peptide bond formation. Anisomycin is one of the strongest activators of ZAKα phosphorylation and the ribotoxic stress response, resulting in rapid p38 phosphorylation (Vind et al., 2020a). We confirmed robust phosphorylation and thus activation of p38 in HEK 293T cells and keratinocytes after anisomycin treatment by immunoblot and flow cytometry with phospho-p38-specific antibodies, suggesting that the ribotoxic stress response is functional in HEK 293T cells and N/TERT-1 keratinocytes (Fig. 4, D, E and F). Anisomycin treatment activated NLRP1 inflammasomes in HEK^NLRP1+ASC^ and N/TERT-1^C1C-EGFP^ cells in a dose-dependent manner (Fig. 4, G and I). The fraction of responding cells was comparable to talabostat in HEK-based systems and substantially stronger than talabostat in keratinocytes, with more than 15% of the cells assembling ASC specks. Robust inflammasome assembly was already observed after 6 h (movies S1-3, fig. S4, F and I). Subsequent stimulation experiments were thus performed for 6 and 20 h (fig. S4, G and J), to allow shorter treatments with toxic inhibitors (6 h) or full comparability to other, slower NLRP1 activators such as talabostat (20 h). ASC speck assembly depended on NLRP1, as neither HEK^NLRP3+ASC^ cells (Fig. 4G), nor NLRP1 or ASC knockout cell lines of N/TERT-1^C1C-EGFP^ assembled specks in response to anisomycin (Fig. 4K). Anisomycin-triggered inflammasome assembly was sensitive to inhibitors of p38α/β or pan-p38 inhibitors (Fig. 4, G and K, Fig. 5, A, C, E, and F). Note that p38 inhibitors do not necessarily inhibit phosphorylation of p38 by upstream kinases, although reduction was observed that may be attributed to feedback loops. As expected, anisomycin treatment also induced the release of IL-1β in a p38-dependent manner (fig. S4, L-N). Primary keratinocytes also responded to anisomycin treatment by robust assembly of ASC specks detected with C1C-EGFP, which were likewise sensitive to p38-inhibitors (Fig. 4L).

**Fig. 5.**
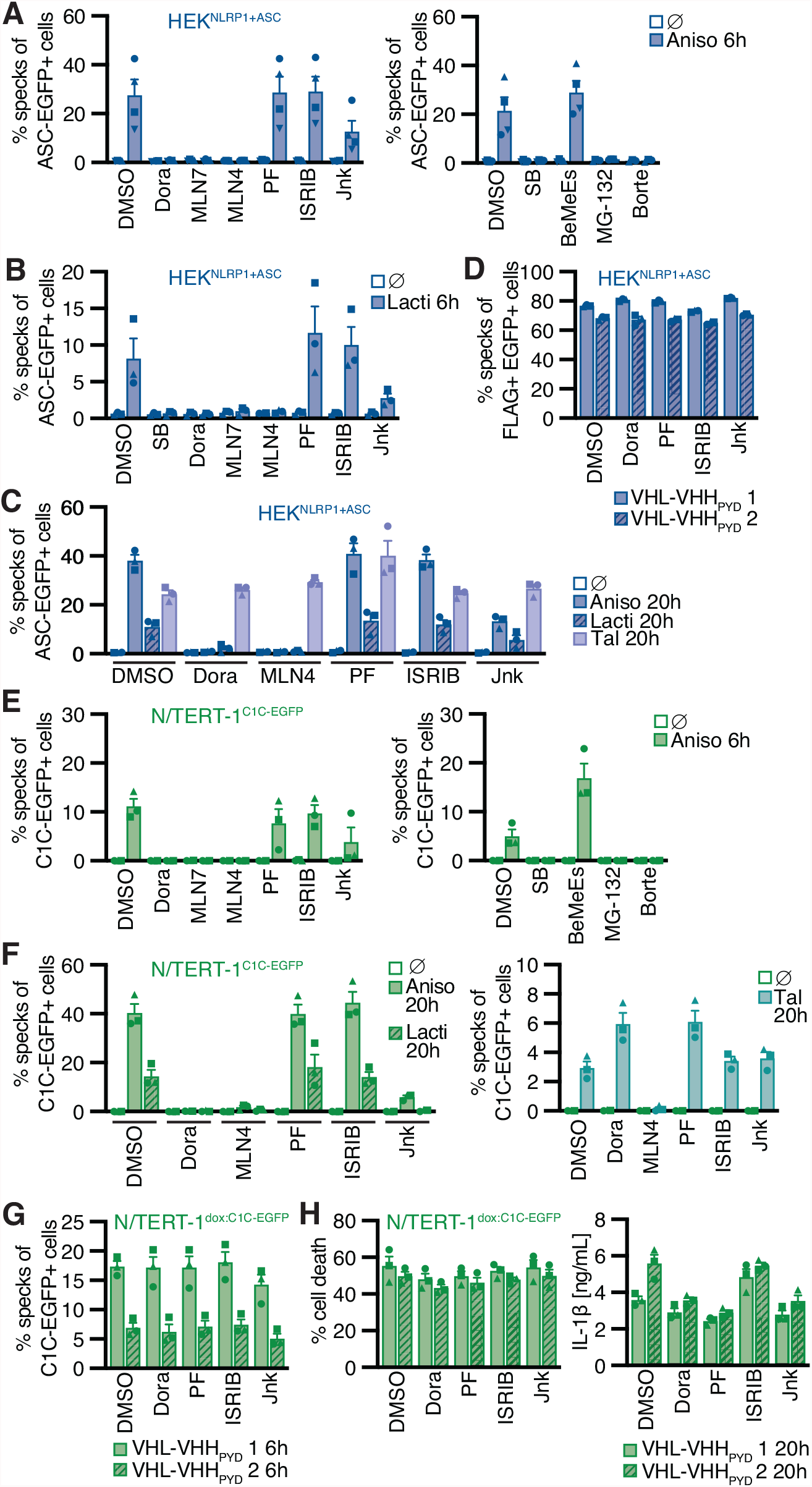
NLRP1 activation by ribotoxic stress response relies on the ubiquitination machinery and proteasomes. HEK^NLRP1+ASC^ (A-D), N/TERT-1^C1C-EGFP^ (E-F), or N/TERT-1 cells inducibly expressing C1C-EGFP and the indicated VHL-VHH fusions (G-H) were stimulated with 15 µM anisomycin (A, C, E, F), 2 µM lactimidomycin (B, F), 30 µM talabostat (C, F), transient (D) or inducible (G-H) expression of VHL-VHH fusions for the indicated time. Cells were stimulated in the presence of 10 µM Dora, 1 µM MLN7, 1 µM MLN4, 1 µM PF, 200 nM ISRIB, 3 µM Jnk-In-8 (Jnk), 20 µM SB, 20 µM Bestatin Methylester (BeMeEs), 1 µM MG-132, 1 µM bortezomib (Borte), or DMSO as indicated. N/TERT-1 cells were always stimulated in the presence 100 µM VX for all flow cytometry experiments. Specking cells were quantified by flow cytometry. Where indicated, cell death was quantified by LDH release and IL-1β release by HTRF. Data represents average values (with individual data points) from three or four independent experiments ± SEM.

Lactimidomycin (Lacti) from *Streptomyces amphibiosporus* and cycloheximide (CHX) inhibit ribosomes by interfering with polypeptide translocation and thus activate the ribotoxic stress response by a different mechanism than anisomycin. Lactimidomycin was found to be a stronger activator of the ribotoxic stress response than CHX (Vind et al., 2020a). Both lactimidomycin and CHX activated NLRP1 inflammasomes in a dose-dependent manner in HEK cells and keratinocytes (Fig. 4, H and J, fig. S4, H and K, movie S4), with a stronger response to lactimidomycin. Concordant with speck responses, IL-1β secretion after CHX treatment was weaker than after anisomycin treatment (fig. S4M). Speck assembly after lactimidomycin stimulation was dependent on NLRP1 and ASC (Fig. 4K). Responses were sensitive to p38 inhibitors, confirming the involvement of the ribotoxic stress response (Fig. 4K, Fig. 5, B, C and F, fig. S4H). Lactimidomycin-induced p38-dependent inflammasome activation was also observed in primary keratinocytes (Fig. 4L).

In the course of our efforts to generate NLRP1 reporter cell lines, we had observed that Blasticidin S (Blasti) also induces low level of NLRP1 inflammasome assembly. Blasticidin impairs termination of translation at the ribosome. We found that blasticidin induces very low level NLRP1 inflammasome assembly at the highest dose in HEK cells and keratinocytes (fig. S4, H and K), but did not initiate any IL-1β response (fig. S4M). ASC speck assembly was sensitive to p38 inhibitors. Weak inflammasome responses in reconstituted HEK cells were difficult to evaluate with regards to quantity and specificity due to the higher background of ectopically expressed ASC-EGFP in untreated HEK^NLRP1+ASC^ cells. Yet, the N/TERT-1^C1C-EGFP^ speck data with virtually no background illustrates the increased sensitivity and specificity of NLRP1 inflammasome detection in this experimental setup, allowing assessment of even the weakest activators.

To evaluate the role of the different p38 isoforms in the activation of NLRP1 inflammasomes, we generated knockout cell lines of p38α and p38β in the background of N/TERT-1^C1C-EGFP^ and compared these to polyclonal ASC knockouts (to compare the maximal inhibition to be expected). Knockout of p38α inhibited anisomycin- and lactimidomycin-induced NLRP1 speck assembly to similar levels as ASC knockouts, while knockouts of p38β did not impair inflammasome assembly (Fig. 4K, upper panels). This suggests that NLRP1 inflammasome assembly is predominantly mediated by p38α in keratinocytes, presumably due to the higher abundance of p38α. In line with these findings, p38 signals detected by immunoblot with a p38 antibody recognizing p38α, β, and γ were substantially diminished in p38α knockout keratinocytes, suggesting that p38α is the most abundant isoform in keratinocytes (fig. S4O). NLRP1 inflammasome assembly triggered by talabostat treatment was not impaired in p38α knockouts and was not sensitive to p38 inhibitors (Fig. 4K, lower panels). In fact, inhibition of p38 substantially boosted the response to talabostat. This suggests that activation of NLRP1 by the ribotoxic stress response follows a mechanism independent of talabostat-mediated inhibition of DPP9 autoinhibition.

Human NLRP1 substantially differs from murine Nlrp1b alleles in the N-terminal PYD and the subsequent linker, which are not found in mNRLP1b (Sastalla et al., 2013). Human NLRP1 is expressed in keratinocytes, while murine keratinocytes do not express Nlrp1b (Sand et al., 2018). To test if the ribotoxic stress response activates both human and murine NLRP1(b), we transiently expressed human NLRP1 or the different mNlrp1b alleles in HEK^ASC^ cells, exploiting that all variants can interact with and activate human ASC. Cells were left untreated or treated with talabostat or anisomycin, followed by quantification of ASC speck responses by flow cytometry. While the background, i.e. the fraction of cells assembling ASC specks merely due to the overexpression of NLRP1(b), was different for each allele, all variants with the exception of mNlrp1b3 were activated by talabostat treatment as described (Gai et al., 2019) (Fig. 4M). While cells expressing human NLRP1 assembled ASC specks in response to anisomycin treatment, none of the murine alleles initiated ASC specks upon anisomycin treatment beyond background levels. To determine which domain of human NLRP1 allowed responsiveness to anisomycin, we constructed an expression vector for a chimeric version of NLRP1 composed of the PYD and linker of human NLRP1 (aa 1-327) and the NACHT-LRR-FIIND-CARD of mNlrp1b (aa 126-1233). Remarkably, cells expressing chimeric h-mNLRP1(b) were able to assemble ASC specks after both talabostat and anisomycin treatment, suggesting that the N-terminal PYD and linker region determine responsiveness to anisomycin.

Many known activators of human NLRP1 and murine Nlrp1b require activity of the proteasome (Bauernfried et al., 2020; Gai et al., 2019; Johnson et al., 2018; Sandstrom et al., 2019). Other cellular factors such as components of the N-end rule pathway are only required for distinct triggers, such as anthrax toxin lethal factor as an activator of Nlrp1b, but not for activation by talabostat (Chui et al., 2019). To define the cellular activities required for NLRP1 activation by the ribotoxic stress response, we quantified inflammasome responses in the presence of different inhibitors in HEK^NLRP1+ASC^ and N/TERT-1^C1C-EGFP^ cells (Fig 5). Activation of NLRP1 with anisomycin, lactimidomycin, CHX, and blasticidin was consistently inhibited by p38 inhibitors SB and Dora (Fig 4G; Fig 5, A-C, E-F; fig S5, A and B). Likewise, NLRP1 activation by the ribotoxic stress response was consistently inhibited by NEDD8 inhibitors. As anisomycin- and lactimidomycin-mediated NLRP1 activation was robustly detected after 6 h, we could also test the effect of an inhibitor of E1 activating enzyme, MLN7, as well as the proteasome inhibitors MG-132 and bortezomib (Borte), which were too toxic for longer experiments. All three inhibitors shut down NLRP1 inflammasome activation completely. Inhibition of MAPKAP2 (PF), the integrated stress response (ISRIB), or the N-end rule pathway (BeMeEs) did not inhibit NLRP1 activation by either activator. Inhibition of JNK kinases (Jnk) only partially affected NLRP1 activation. Importantly, inhibition of p38, neddylation, E1, or the proteasome for the same duration of time did not inhibit NLRP3 activation by LPS and nigericin in HEK^NLRP3+ASC^ (fig, S5, C and D); E1 and proteasome inhibitors even enhanced NLRP3 responses. This indicates that the requirement for these cellular activities is specific for NLRP1, and not inflammasomes in general. Activation of NLRP1 by talabostat (Fig. 5, C and F) or by ubiquitination of the NLRP1 N-terminus (Fig. 5, D, G and H) did not require p38 activity, suggesting that p38 activation is involved in a signaling cascade upstream of NLRP1, but not required for NLRP1 function, priming, or licensing in general. Transient expression of VHL-VHH constructs in HEK^NLRP1+ASC^ or C1C-EGFP in N/TERT keratinocytes was not inhibited by either of the drugs, ruling out any indirect effects of gene expression (fig. S5, E and F).

Activation of NLRP1 by ATP depletion or H_2_O_2_ in HEK^NLRP1+ASC^ was not sensitive to inhibitors of p38 activity, E1 enzymes, neddylation, or the proteasome, suggesting yet another mechanism of activation independent of the ribotoxic stress response (fig. S5G).

Taken together, we find that diverse activators of the ribotoxic stress response activate human NLRP1. This response is distinct from NLRP1 activation by inhibition of DPP9 as it depends on p38 kinase activity and requires features found in the N-terminus of human NLRP1, but not murine Nlrp1b alleles. As several of the described NLRP1 activators efficiently inhibit translation, NLRP1 activation does not depend on p38-dependent gene expression, but rather on the kinase activity of p38 directly. As described for other NLRP1 stimuli, NLRP1 activation through the ribotoxic stress response required E1 activity, neddylation, and proteasome activity, indicating that these cellular functions are required for most mechanisms of NLRP1 inflammasome activation (with the notable exception of ATP depletion and ROS). The ubiquitin-proteasome pathway likely contributes to the N-terminal degradation of NLRP1 itself.

### Alphavirus-induced NLRP1 activation is also dependent on p38 kinase activity

Infection of human keratinocytes with the model alphavirus SFV as well as cytosolic delivery of dsRNA was reported to activate NLRP1 inflammasomes by direct binding of dsRNA to NLRP1 (Bauernfried et al., 2020). To study NLRP1 activation in response to viral infection, we thus infected HEK^NLRP1+ASC^, HEK^NLRP3+ASC^, and N/TERT-1^C1C-EGFP^ cells with SFV, the closely related Sindbis virus (SINV), as well as vesicular stomatitis virus (VSV), a negative-sense single-stranded RNA virus. We quantified infection by staining for dsRNA (SFV, SINV) or the VSV-G protein, and only included infected cells in the flow cytometry analysis of inflammasome activation. Nearly all of the cells were infected with the respective viruses, but only SFV and SINV induced a robust assembly of ASC specks in HEK^NLRP1+ASC^ and N/TERT-1^C1C-EGFP^ cells 20 h post infection (Fig. 6, A and B). The response in the HEK-based system was weak compared to talabostat, while keratinocytes exhibited robust speck formation. SFV was a stronger inflammasome activator than SINV in both reporter cell lines and the formation of specks in SFV-infected N/TERT-1^C1C-EGFP^ was not detectable in ASC and NLRP1 knockout cells (Fig. 6C). Treatment of the cells with the V-ATPase inhibitor bafilomycin A1 (BafA) completely abolished alphavirus infection, as alphaviruses require endosomal acidification to fuse with the limiting membrane of endosomes (Marsh et al., 1983) (fig. S6, A and B). The treatment also abrogated ASC speck formation, demonstrating that cytosolic delivery of viral genomes was critical to activate NLRP1 in the reporter cells. SFV- and SINV-infected N/TERT-1^C1C-EGFP^ cells released IL-1β in a caspase-1-dependent manner, which was also inhibited by BafA (fig. S6C).

**Fig. 6.**
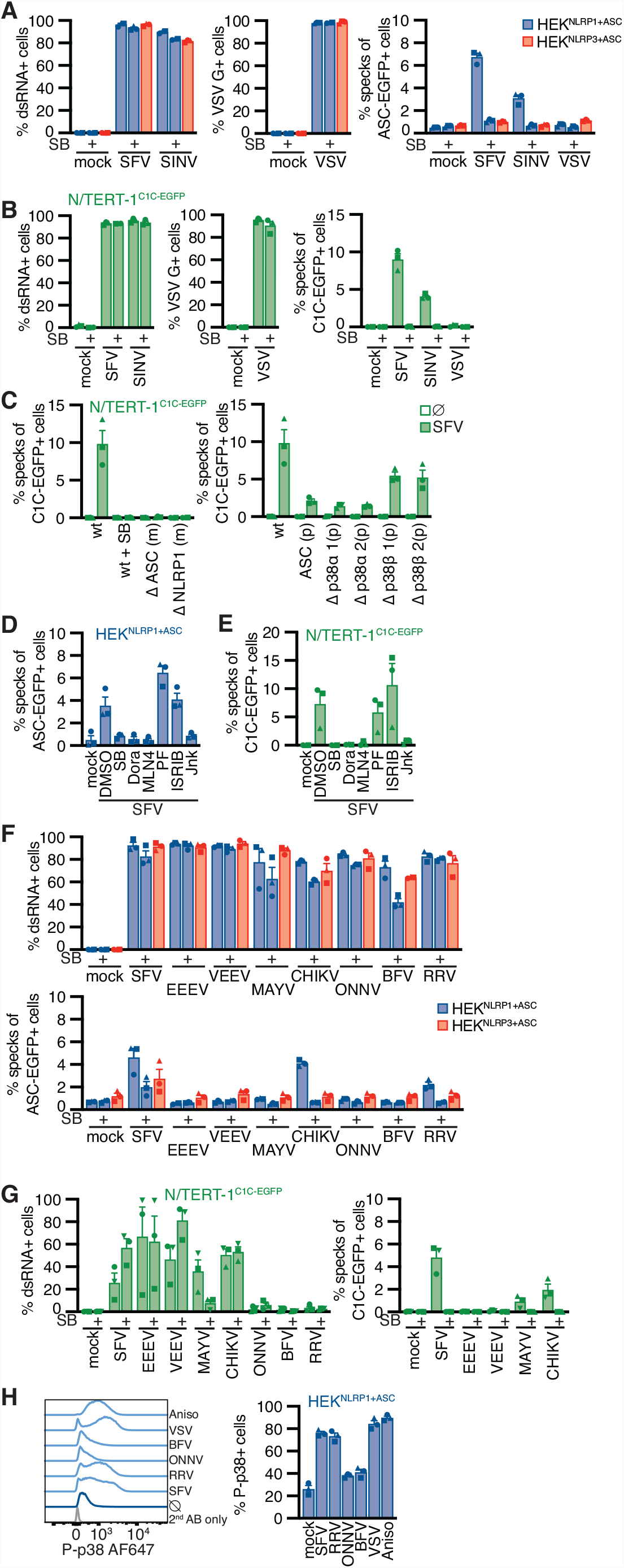
Alphavirus infection activates NLRP1 in a p38-dependent manner. (**A-C**) HEK^NLRP1+ASC^ and HEK^NLRP3+ASC^ (A), N/TERT-1^C1C-EGFP^ (B), or knockout strains derived from N/TERT-1^C1C-EGFP^ (C) were infected with Semliki Forest virus (SFV), Sindbis virus (SINV), or vesicular stomatitis virus (VSV) at a multiplicity of infection (MOI) of 1 (HEK) or 5 (N/TERT-1) for 20 h in the presence or absence of 20 µM SB. Infected cells were stained with antibodies for double-stranded RNA (dsRNA) (SFV, SINV), or VSV G (VSV) and infection and speck assembly in infected cells was quantified by flow cytometry. (**D, E**) HEK^NLRP1+ASC^ (D), or N/TERT-1^C1C- EGFP^ (E) cells were infected with SFV as in (A), cultivated for a 20 h in the presence of the indicated drugs, and analyzed for specks as in (A). (**F, G**) HEK^NLRP1+ASC^ (F), or N/TERT-1^C1C-EGFP^ (G) cells were infected with SFV (F: MOI 1/G: MOI 5), Eastern equine encephalitis virus (EEEV) (MOI 1/5), Venezuelan equine encephalitis virus (VEEV) (MOI 1/50), Chikungunya virus (CHIKV) (MOI 1/50), Mayaro virus (MAYV) (MOI 25/50), o’nyong nyong virus (ONNV) (MOI 25/50), Barmah Forest Virus (BFV) (MOI 25/50) or Ross River virus (RRV) (MOI 25/50) for 20 h in the absence or presence of 20 µM SB. Infection and speck responses were quantified as in (A). (**H**) HEK^NLRP1+ASC^ cells were infected with the indicated viruses as in (A) and (F) and stained for phospho-p38. Control cells were treated with 15 µM anisomycin. Representative histograms (left), and average fractions of P-p38-positive cells from three independent experiments (right) are displayed. N/TERT-1 cells were always infected in the presence 100 µM VX for all flow cytometry experiments. Data represents average values (with individual data points) from three independent experiments ± SEM.

Since we had found that activators of the ribotoxic stress response activate human NLRP1 in a p38-dependent manner, we wondered whether the activation by alphaviruses also depends on p38 kinase activity. ASC speck formation in HEK^NLRP1+ASC^ and N/TERT-1^C1C-EGFP^ cells infected with SFV and SINV was completely inhibited by the p38α/β inhibitor SB (Fig. 6, A, B, and C). In line with that, IL-1β release by alphavirus-infected keratinocytes was reduced by SB (fig. S6C). Polyclonal knockouts of p38α in N/TERT-1^C1C-EGFP^ cells inhibited SFV-induced NLRP1 speck assembly to similar levels as ASC knockouts (Fig. 6C), while infection was not altered in the knockout cells (fig. S6D). The pan-p38 inhibitor Dora, the neddylation inhibitor MLN4, and the JNK inhibitor Jnk completely inhibited inflammasome assembly in HEK^NLRP1+ASC^ and N/TERT-1^C1C-EGFP^ cells after infection with SFV (Fig. 6, D and E). MLN4 reduced infection of keratinocytes, but the fraction of infected cells was sufficient for analysis (fig S6, E and F). Like NLRP1 activation by triggers of the ribotoxic stress response, we thus found that NLRP1 inflammasome activation induced by alphavirus infection depends on p38 kinase activity and neddylation.

Double-stranded RNA intermediates of SFV have been proposed to be the relevant trigger for NLRP1 (Bauernfried et al., 2020), and we also observed that N/TERT-1 keratinocytes assemble inflammasomes upon transfection with poly(I:C) (Fig. 2I). We thus tested whether activation of NLRP1 inflammasome in N/TERT-1^C1C-EGFP^ cells by dsRNA was similarly dependent on p38. We found that both the response to cytosolic dsRNA, as well as the inflammasome response to DNA in the reporter keratinocytes was completely inhibited by the p38α/β inhibitor SB (fig. S6G).

We next tested whether NLRP1 is also important in the detection of alphaviruses that are clinically relevant as human pathogens. We thus infected the HEK and keratinocyte reporter cells with a panel of human pathogenic alphaviruses including Eastern equine encephalitis virus (EEEV), Venezuelan equine encephalitis virus (VEEV), Mayaro virus (MAYV), Chikungunya virus (CHIKV), o’nyong-nyong virus (ONNV), Barmah Forest virus (BFV), and Ross River virus (RRV) in the presence and absence of SB. All tested viruses were able to infect both HEK^NLRP1+ASC^ and HEK^NLRP3+ASC^ cells, as confirmed by staining of dsRNA (Fig. 6F). In addition to the positive control SFV, we found that CHIKV and RRV activated NLRP1-dependent inflammasomes. ASC assembly in response to all three alphaviruses was inhibited by the p38α/β inhibitor SB. We were not able to infect the N/TERT-1^C1C-EGFP^ cells with ONNV, BFV, and RRV (Fig. 6G) and the fraction of infected cells was generally more variable for keratinocytes. However, analysis of inflammasome assembly in infected cells revealed clear ASC specking responses in cells infected with SFV, MAYV, and CHIKV. We cannot rule out that the weak NLRP1 activation in response to MAYV infection was not detected in HEK^NLRP1+ASC^ cells due to the higher background caused by ectopically expressed ASC-EGFP. CHIKV-induced ASC speck assembly in the reporter keratinocytes was also dependent on p38 activity. Since SB strongly decreased MAYV infection, we could not assess whether the weak inflammasome response depends on p38.

It is likely that related alphaviruses expose similar viral molecules in the cytosol and that they hijack similar cellular functions. Yet, only distinct alphaviruses triggered NLRP1 inflammasome assembly. We thus wondered if differential NLRP1 activation correlated to differences in p38 activation. We infected HEK^NLRP1+ASC^ cells with two alphaviruses shown to trigger ASC specks (SFV and RRV) as well as two viruses that did not initiate inflammasome assembly (ONNV and BFV) (fig. S6H). We detected robust phosphorylation of p38 by flow cytometry in cells infected with SFV and RRV that was comparable to p38 phosphorylation in anisomycin-treated cells (Fig. 6H). Cells infected with ONNV and BFV, however, did not trigger p38 phosphorylation (Fig. 6H). The ability to activate p38 thus correlated with NLRP1 activation, indicating that MAP kinase signaling is the critical determinant of NLRP1 activation by alphaviruses. Surprisingly, infection with the control virus VSV induced p38 phosphorylation as well, indicating that either p38 activation by alphaviruses differs from activation by VSV, or that VSV counteracts p38-dependent NLRP1 inflammasome stimulation.

Activation of the ribotoxic stress response and p38 phosphorylation relies on the α splice variant of MAP3 kinase ZAK (ZAKα). To test whether NLRP1 activation by alphaviruses engages a similar signaling cascade, we generated knockouts of ZAK as well as a control MAP3 kinase, TAO kinase 2 (TAOK2), in the background of N/TERT-1^C1C-EGFP^ cells (fig. S7A). As expected, NLRP1 activation by talabostat was not affected by either of the knockouts (Fig. 7A). Anisomycin-induced ASC speck assembly was completely abrogated in ZAK-knockout cells, validating the critical role of ZAKα for the ribotoxic stress response and the resulting p38-dependent NLRP1 activation. In comparison, SFV-induced NLRP1 inflammasome activation was reduced by around 70% in ZAK-knockout cells, but a fraction of cells was still responsive, suggesting that p38 activation can be mediated by other MAP3 kinases in the context of alphavirus infections. All knockout cells were properly infected as quantified by dsRNA staining (fig. S7B).

**Fig. 7.**
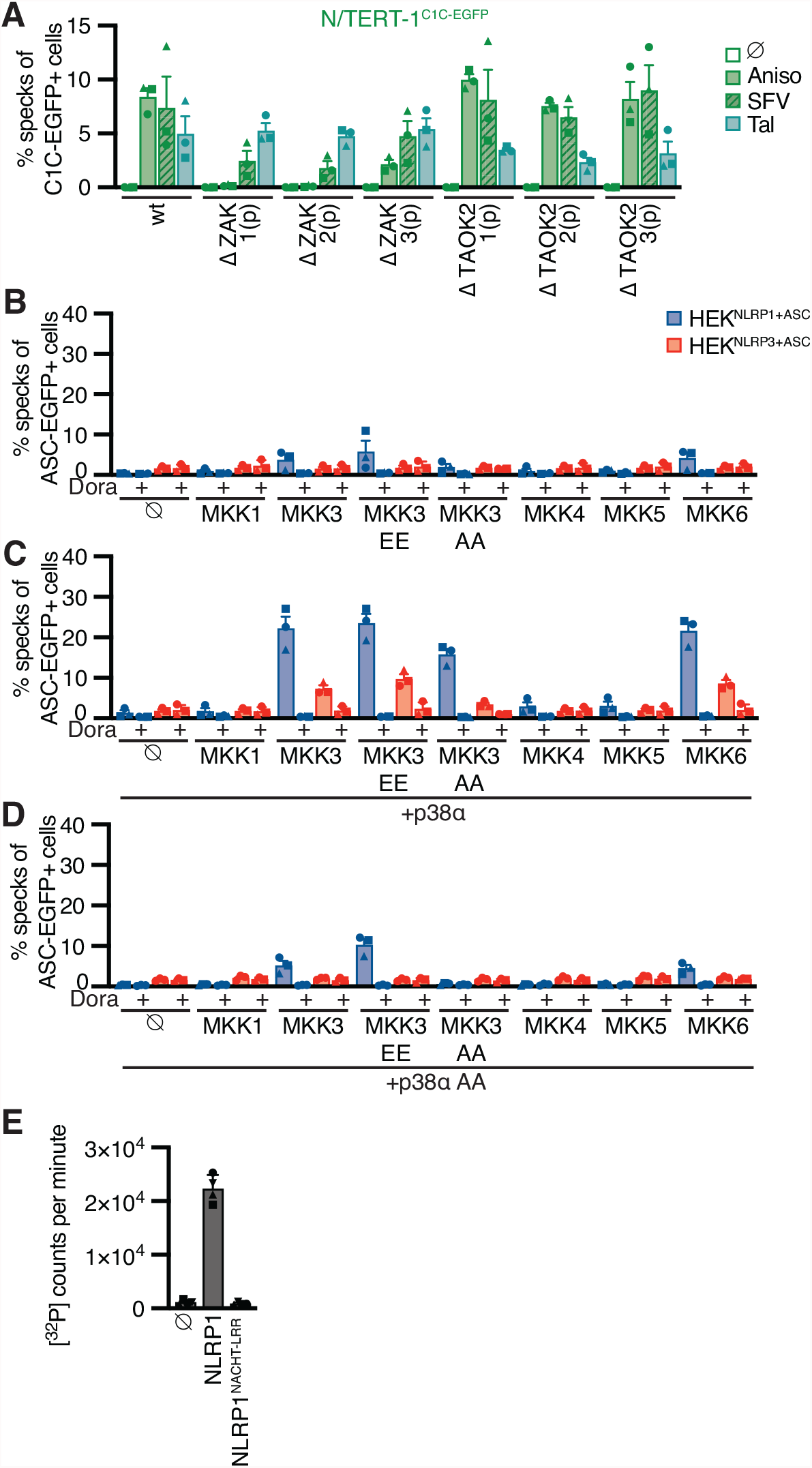
P38 activation is sufficient for NLRP1 activation. (**A**) N/TERT-1^C1C-EGFP^ wt or knockout strains derived from N/TERT-1^C1C-EGFP^ were treated with 15 µM anisomycin, 30 µM talabostat, or infected with SFV at an MOI of 5 for 20 h in the presence or absence of 20 µM SB. Speck assembly was quantified by flow cytometry, in case of SFV limited to infected cells (see fig. S7A). (**B-D**) HEK^NLRP1+ASC^ or HEK^NLRP3+ASC^ cells were transiently transfected with empty vectors (B), expression vectors for p38α (C) or p38α T180A Y182A (p38α AA) (D) in combination with expression vectors for MKK1, MKK3, MKK3 S218E T222E (MKK3 EE), MKK3 S218A T222A (MKK3 AA), MKK4, MKK5, or MKK6. Where indicated, cells were treated with 20 µM Dora. Cells with ASC-EGFP specks were quantified by flow cytometry. Data represents average values (with individual data points) from three independent experiments ± SEM. (**E**) 0.2 µM recombinant p38d was incubated with recombinant 7 µM MBP-NLRP1 or MBP-NLRP1^ΔPYD-linker^ (aa 230-990) or no substrate at 30 °C for 30 min in kinase buffer containing [^32^P]-γ-ATP. Protein-associated ^32^P was quantified by liquid scintillation counting. Data represents average values (with individual data points) from four independent experiments ± SD.

Taken together, we recapitulated NLRP1 activation by SFV infection in NLRP1 HEK and keratinocyte reporter cells. We identified four additional alphaviruses, SINV, RRV, MAYV, and CHIKV, that activate NLRP1, supporting the relevance of NLRP1 as a sensor for alphavirus infection. Interestingly, NLRP1 activation was not universally observed for all alphaviruses tested. Since alphavirus-induced ASC speck formation was completely dependent on p38 kinase activity, we identified activation of p38 as a key event for NLRP1 activation.

### P38 activity is sufficient for NLRP1 activation

Having shown that p38 activity was required for NLRP1 activation by the ribotoxic stress response and alphavirus infection, we next assessed whether p38 activation itself was sufficient for NLRP1 activation. We transiently overexpressed the MAP2 kinases MKK1, MKK3, MKK4, MKK5, and MKK6 in HEK^NLRP1+ASC^ and HEK^NLRP3+ASC^ cells. We also included phosphomimetic glutamate and inhibitory alanine mutants of S218 and T222 in the activation loop of MKK3, yielding MKK3 EE or MKK3 AA, respectively. Overexpression of MKK3, MKK3 EE, or MKK6, the upstream kinases that directly phosphorylate p38, indeed initiated weak NLRP1 inflammasome activation (Fig. 7B). Activation was sensitive to the pan-p38 inhibitor Dora, confirming that the observed inflammasome assembly indeed relied on p38 activity. Importantly, the related MAP2 kinases MKK1, MKK4, and MKK5, which phosphorylate other MAP kinases but not p38, did not initiate inflammasome assembly in HEK^NLRP1+ASC^. Overexpression of none of the MAP2 kinases activated NLRP3 inflammasome assembly in HEK^NLRP3+ASC^. Experiments to artificially trigger p38 activity typically rely on the combined ectopic expression of p38 isoforms and their upstream kinases (Raingeaud et al., 1996). In line with these findings, overexpression of p38α (Fig. 7C), p38β, p38γ, or p38δ (fig. S7, C-E) alone did not initiate NLRP1 inflammasome assembly. However, combined expression of p38α with MKK3, MKK3 EE, or MKK6 triggered robust NLRP1 inflammasome assembly (Fig. 7C). Interestingly, MKK3 AA bearing a mutated activation loop that cannot be phosphorylated still enhanced p38α inflammasome assembly. None of the other MAP2 kinases stimulated p38α-mediated NLRP1 activation. Dora treatment abrogated all observed NLRP1 inflammasome assembly, indicating that the observed ASC specks genuinely relied on p38 activity. Co-expression of MAP2 kinases with p38α bearing a mutated activation loop (p38α T180A Y182A) did not enhance NLRP1 activation beyond the levels observed upon overexpression of upstream kinases alone (Fig. 7D). Remarkably, p38α activation by overexpression also induced some level of NLRP3 activation, albeit to substantially lower levels than observed for NLRP1. Similar NLRP1 activation was observed when p38β, p38γ, or p38d were co-expressed with MAP2 kinases (fig. S7, C-E). NLRP1 activation was again sensitive to inhibition by pan-p38 inhibitor Dora. In line with the higher IC_50_ values of this drug for p38δ (Kuma et al., 2005), inhibition was incomplete for this isoform.

NLRP1 activation by targeted ubiquitination of the PYD did not require p38 activity, and neither did the activation of NLRP1 by talabostat. This indicates p38 is not required for NLRP1 activation per se, but that p38 contributes to an upstream signaling cascade that integrates information from different stress signaling cascades to activate NLRP1. We could show that p38 activation was sufficient for NLRP1 inflammasome activation, and that activation was likely not a consequence of p38-dependent gene expression. To test whether p38 is able to phosphorylate NLRP1 directly, we performed radiometric in vitro kinase assays with recombinant p38δ and purified full length NLRP1 expressed in insect cells as fusion to maltose-binding protein (MBP-NLRP1, fig. S7F). We found that MBP-NLRP1 could be robustly phosphorylated by p38δ, as detected by transfer of radioactive phosphate from [^32^P]-γ-ATP to purified NLRP1 protein (Fig 7E). Importantly, MBP-NLRP1^aa 230-990^, comprising part of the linker, NACHT, and LRR of NLRP1, could not be phosphorylated by p38δ. This indicates that either the N-terminal 230 amino acids or the C-terminal FIIND-CARD of NLRP1 are the substrate of p38. Given that the N-terminal 327 amino acids of human NLRP1 were sufficient to render murine Nlrp1b responsive to anisomycin, phosphorylation of NLRP1 by p38 likely occurs in the N-terminal PYD or linker of NLRP1.

In sum, we find that p38 can directly phosphorylate NLRP1 and that p38 activation in reconstituted HEK^NLRP1+ASC^ is sufficient to initiate the assembly of NLRP1 inflammasomes.

## Discussion

As the largest organ, the human skin is exposed to a variety of physico-chemical and biological insults, including UV irradiation and infection with arthropod-borne viruses. These can trigger the assembly of NLRP1 inflammasomes in keratinocytes (Fenini et al., 2020). By using reporters for ASC speck assembly rather than cytokine measurements alone, we could robustly quantify inflammasome assembly in keratinocytes, even though the fraction for responding cells is lower than observed for myeloid cells. This may avoid the massive cell death of abundant keratinocytes in response to threats that can be locally controlled.

In this study, we find that a number of different stress signals, including UVB irradiation, inhibition of translation by antibiotics, as well as infection with arthropod-borne alphaviruses, including the relevant human pathogen CHIKV, converge on the activation of p38 MAP kinase signaling. P38 kinase activity lastly activates NLRP1 inflammasomes by direct phosphorylation. Importantly, we find that strong p38 activation is sufficient to activate NLRP1, suggesting that p38 signaling acts as a rheostat integrating different stress responses to ultimately trigger irreversible NLRP1 inflammasome assembly if the critical threshold is overcome.

The role of keratinocytes in the inflammatory response to UV and other triggers is well established. Extending previous studies on the UV-induced ribotoxic stress response (Iordanov et al., 1998; Wang et al., 2005), and the underlying MAP kinase signaling cascade in epithelial cells (Vind et al., 2020b, 2020a), as well as the role of p38 in inflammation (Fenini et al., 2018b), we conclude the following model: UV-induced damage of ribosomal RNA causes ribosome clashing and activation of the ribotoxic stress response in keratinocytes. The same pathway is initiated by interfering with ribosome function, such as inhibition of peptide bond formation or peptide translocation by anisomycin or lactimidomycin, respectively. Other toxins that activate the ribotoxic stress response, such as ricin and diphtheria toxin, likely stimulate the same pathway. Ribotoxic stress responses trigger a phosphorylation cascade entailing the activation of ZAKα (MAP3K), MKK3/MKK6 (MAP2K), and finally p38 (MAPK). We find that p38α plays a predominant role in the activation of NLRP1 inflammasomes in keratinocytes. Yet, ectopic activation of any p38 isoform by overexpression of its upstream kinases MKK3 or MKK6 is sufficient to initiate NLRP1 inflammasome assembly. P38 can directly phosphorylate the N-terminus of NLRP1 and thereby triggers its activation in a pathway that relies on neddylation, ubiquitin activation, and proteasome activity. Murine Nlrp1b alleles are not responsive to anisomycin, but the capability to be activated by anisomycin can be transferred with the human NLRP1 N-terminus (PYD + linker). It is likely that phosphorylation of unique sites of human NLRP1 allows the ubiquitination of the NLRP1 N-terminus by NEDD8-dependent cullin ligases, followed by N-terminal degradation, recruitment of ASC by NLRP1^UPA^-CARD, as well as assembly of ASC specks. ASC specks, in turn, are the sites of caspase-1 activation, resulting in IL-1β maturation and GSDMD-dependent cell death by pyroptosis.

Alphaviruses are transmitted by mosquito bites and likely first replicate in keratinocytes and fibroblasts in the skin before infection is spread by migratory dendritic cells. A sensitive response system in the skin is therefore beneficial to detect and contain infecting alphaviruses before any systemic spread can occur. Viral infection may even be detected in non-permissive cells, in which no full replication occurs. In line with this, full CHIKV replication in keratinocytes is severely impaired (Bernard et al., 2015). It is likely that alphaviruses, in turn, have evolved mechanisms to escape detection by NLRP1. Remarkably, we find that NLRP1 is exclusively activated by those alphaviruses that mediate strong p38 activation. As viral molecules in the cytosol are similar between different alphaviruses, it is possible that some alphaviruses can control p38 activation in the host and thus prevent NLRP1 activation. That VSV activates p38, but not NLRP1 inflammasomes, suggests that viruses can either interfere with inflammasome assembly downstream of p38, or that infection can initiate different qualities of MAP kinase responses that are not sufficient for NLRP1 activation and require further study.

Remarkably, viral NLRP1 activation does not seem to exclusively rely on ZAKα, suggesting that different MAP3 kinases can feed into the same activation mechanism, highlighting the role of p38 as a central signaling hub that integrates information from various inputs. While we found that some kinase inhibitors specifically inhibit viral activation of NLRP1 (data not shown), the corresponding additional kinase(s) still await identification. Cytosolic delivery of dsRNA, as well as DNA, is sufficient to activate NLRP1 in a p38-dependent manner in keratinocytes, but not in reporter HEK cells. This suggests that nucleic acids contribute to p38 stimulation, but that other cues may be required to activate NLRP1. Such additional signals may be provided by infection, as SFV infection was able to activate NLRP1 in both keratinocytes and HEK cells with reconstituted NLRP inflammasomes.

As a first line of defense, the skin encounters a variety of biotic and abiotic stresses that have to be interpreted to elicit inflammation with highest sensitivity, while at the same time limiting unnecessary collateral damage. We find that p38 MAP kinase signaling integrates different stress inputs and can – as a result of sufficient activation – directly phosphorylate NLRP1 to trigger its activation. While p38 is ubiquitously expressed throughout the body, NLRP1 expression is limited to few distinct cell types. In keratinocytes, p38 signaling can thus be co-opted to feed into a more drastic (and wide-spread) inflammatory response by activating NLRP1.

## Methods

### Cell lines

Human embryonic kidney (HEK) 293T cells (ATCC), and Syrian baby hamster kidney (BHK)-21 cell clone BSR-T7/5 (*Mesocricetus auratus*, a kind gift of SeanWhelan, Harvard Medical School) were cultivated in DMEM containing 10% FBS and GlutaMax; BHK-21/J cells (*Mesocricetus auratus*, a kind gift of Charles M. Rice, Rockefeller University) were cultivated in MEM containing 7.5% FBS, 1% nonessential amino acids, and 1 % L-glutamine (all Thermo Fisher Scientific). Human N/TERT-1 keratinocytes (a kind gift of James Rheinwald, Harvard Medical School) were cultivated in Keratinocyte serum-free medium (Thermo Fisher Scientific) supplement with 0.5x bovine pituitary extract (BPE), 0.2 mg/mL epidermal growth factor (EGF), 1x penicillin/streptomycin (PS), and 0.3 mM CaCl_2_ (resulting in a final concentration of 0.4 mM CaCl_2_). Primary Normal Human Epidermal Keratinocytes (NHEK) were obtained from PromoCell (Heidelberg, Germany) and cultivated in Keratinocyte Growth Medium 2 (PromoCell). Unless otherwise mentioned, genetically modified cell lines were generated by lentiviral transduction using lentivirus produced with packaging vectors psPax2 and pMD2.G (kind gifts from Didier Trono, École polytechnique fédérale de Lausanne, Switzerland) and antibiotic selection. Lentiviral vectors for constitutive expression under the control of the ubiquitin C promoter (pUbC) were constructed by Gateway cloning (Thermo Fisher Scientific) using vectors modified from pRRL (a kind gift of Susan Lindquist, Whitehead Institute of Biomedical Research). HEK 293T cells expressing ASC-EGFP under control of pUbC, HEK^ASC^, were generated by transduction with a dilution series of lentivirus to achieve single insertions. Individual clones were cultivated and tested for minimal background of ASC-EGFP specks. Derivatives of HEK^ASC^ expressing NLRP1-HA (HEK^NLRP1+ASC^; cell line H8-1), or NLRP3-HA (HEK^NLRP3+ASC^; cell line H98) under control of pUbC from single insertions were generated using the same protocol and clones selected based on optimal signal-to-noise ratio and background. N/TERT-1 expressing caspase-1^CARD^-EGFP (C1C-EGFP) controlled by pUbC (N/TERT-1^C1C-EGFP^, cell line K14), were similarly generated by lentiviral transduction. Knockouts of N/TERT-1^C1C-EGFP^ were generated by lentiviral transductions using vectors modified based on pLenti CRISPR v2 (a kind gift of Feng Zhang, Broad Institute) (see table S1 for sgRNA target sequences). Monoclonal knockouts were generated using limiting dilutions of polyclonal cell lines generated by lentiviral transduction, and selection with antibiotics; best clones were selected based on analysis with OutKnocker (Schmid-Burgk et al., 2014) and immunoblot. Polyclonal knockouts were generated by lentiviral transduction and antibiotic selection and verified by immunoblot. Derivatives of N/TERT-1 expressing C1C-EGFP and VHHs, VHL-VHH fusions, or MAP2 kinases under a bidirectional dox-inducible promoter were generated using lentiviral vector pInducer20bi-NA, a derivative of pInducer20-NA (Schmidt et al., 2016a) using the promoter from pTRE3G-BI (Takara Bio).

**Table S1.**
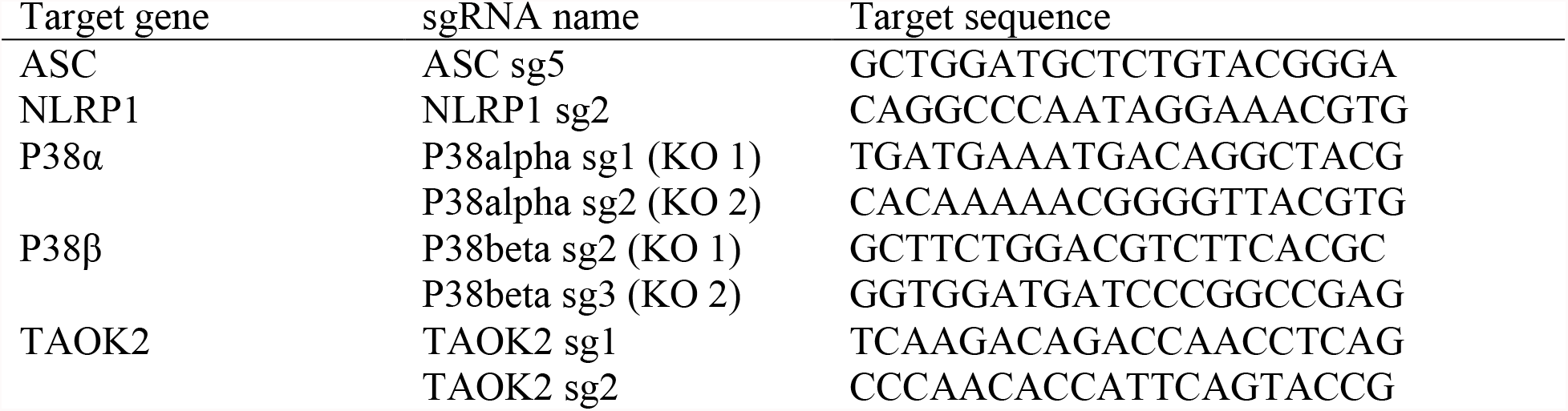
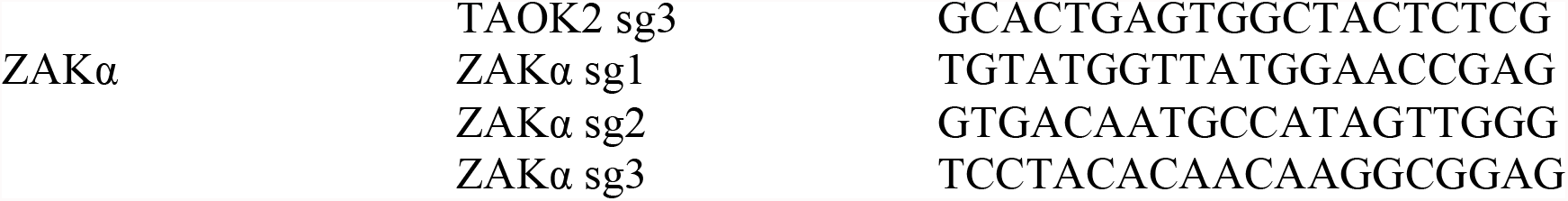
sgRNA target sequences used for the generation of knockout cell lines.

### Viruses

All experiments involving viruses were conducted in respective Biosafety Level 2 or 3 laboratories. Semliki Forrest Virus (SFV) 4 (a kind gift of Giuseppe Balistreri and Ari Helenius, ETH Zurich, Switzerland), Sindbis virus (SINV) strain Toto 1101 (recovered from *in vitro* transcribed RNA (Rice,’ et al., 1987)), and vesicular stomatitis virus (VSV) strain Indiana (recovered from plasmid as described in (Whelan et al., 1995)) were amplified in BSR-T7 cells. Eastern equine encephalitis virus (EEEV), Venezuelan equine encephalitis virus (VEEV), Mayaro virus (MAYV), o’nyong-nyong virus (ONNV), Barmah Forest virus (BFV), and Ross River virus (RRV) (kindly provided by Klaus Grywna (Grywna et al., 2010)) were amplified in BHK-J cells. Chikungunya virus (CHIKV) was derived from an infectious cDNA clone as described earlier (Kümmerer et al., 2012). Virus-containing supernatants clarified from cell debris were aliquoted and frozen at −80°C. Viral titers of SFV, SINV, and VSV were determined by flow cytometry using anti-dsRNA antibody J2 (Schonborn et al., 1991), or anti-VSV G I14 (clone 1E9F9, (Lefrancois and Lyles, 1982)) combined with fluorescent secondary antibodies. Viral titers of EEEV, VEEV, MAYV, ONNV, BFV, RRV, and CHIKV were determined by plaque assay using BHK-J cells (Karliuk et al., 2021).

### Proteins

#### Expression and purification of His-NLRP1^PYD^ and GST-NLRP1^PYD^

Expression vectors for human His-NLRP1^PYD^ (aa 1-92) and GST-NLRP1^PYD^ were generated by Gateway cloning with destination vector pDEST17 (Thermo Fisher Scientific) and a customized destination vector based on pGEX-2T (a kind gift of Mikko Taipale, Whitehead Institute of Biomedical Research). Proteins were expressed in *Escherichia (E*.*) coli* LOBSTR (Andersen et al., 2013) (His-NLRP^PYD^) or *E. coli* BL21 (GST-NLRP1^PYD^) cells induced with 0.2 mM IPTG at an OD600 of 0.6. Cells were cultivated for 20 h at 18° C, and lysed by French Press or sonication. His-NLRP^PYD^ was purified by Ni-NTA affinity chromatography and gel filtration with a HiLoad 16/600 Superdex 75 pg column in buffers containing 50 mM Tris pH 8.0 and 500 mM NaCl. GST-NRLP1^PYD^ was purified by affinity chromatography with glutathione resin and gel filtration with a HiLoad 16/600 Superdex 75 pg column in buffers containing 50 mM Tris pH 8.0 and 500 mM NaCl.

#### Expression and purification of His-MBP-NLRP1

Full length human NLRP1 (aa 1-1473) and NLRP1^NACHT-LRR^ (aa 230-990) constructs were expressed as N-terminal MBP fusion proteins in baculo virus infected *Spodoptera frugiperda* (*Sf9*) insect cells. For expression, 0.5 L of *Sf9* cells at a density of 2.0 × 10^6^ were infected with 5% P2 virus and incubated at 28°C for 48 h before harvesting the cells by centrifugation at 1125 x g for 20 min. Cells were washed once with cold PBS and then lysed by sonication in buffer A (50 mM HEPES pH 7.5, 150 mM NaCl, 0.5 mM TCEP), freshly supplemented with 1 mM PMSF. The lysate was clarified by centrifugation at 75,000 x g for 30 min at 10°C. Proteins were affinity purified using an MBPTrap HP (GE Healthcare) affinity column and eluted in buffer A supplemented with 10 mM maltose.

#### Expression and purification of nanobodies

Nanobody coding sequences were cloned into pHEN6-based bacterial, periplasmic expression vectors with C-terminal HA-His_6_ tags using Gibson cloning (New England Biolabs). Nanobodies were produced in *Escherichia coli* WK6 transformed with the respective expression vectors. Expression cultures were grown in Terrific Broth (TB), and expression was induced with 1 mM IPTG at an OD_600_ of 0.6, followed by cultivation at 30° C for 16 h. Bacterial pellets were resuspended in TES buffer (200 mM Tris-HCl pH 8.0, 0.65 mM EDTA, 0.5 M sucrose), and periplasmic extracts generated by osmotic shock in 0.25x TES, followed by Ni-NTA purification and desalting by PD MiniTrap G-25 columns (GE Healthcare Life Sciences).

### Antibodies

The following antibodies were used: rabbit anti-ASC (AdipoGen Cat# AG-25B-0006, RRID:AB_2490440), mouse anti-GFP clone JL-8 (Takara Bio Cat# 632380, RRID:AB_10013427), rabbit anti-FLAG clone D6W5B (Cell Signaling Technology Cat# 14793, RRID:AB_2572291), mouse anti-HA clone 16B12 (HA.11, Biolegend, Cat# 901503, RRID:AB_2565005), mouse anti-gamma H2A.X (phospho S139) clone 9F3 (Abcam Cat# ab26350, RRID:AB_470861), mouse anti-NLRP1 Clone 9F9B12 (BioLegend Cat# 679802, RRID:AB_2566263), rabbit anti-p38 (Cell Signaling Technology Cat# 9212, RRID:AB_330713), anti-phospho-p38 (T180/Y182) clone D3F9 (Cell Signaling Technology Cat# 4511, RRID:AB_2139682), mouse anti-dsRNA clone J2 (SCICONS Cat# 10010200, RRID:AB_2651015), mouse anti-vinculin clone hVIN-1 (Sigma-Aldrich Cat# V9131, RRID:AB_477629), mouse anti-VSV G I14 clone 1E9F9 (from D.S. Lyles, kindly provided by Ari Helenius).

### Small compound inhibitors

The following small compound inhibitors were used: anisomycin (Sigma-Aldrich), bafilomycin A1 (Sigma-Aldrich), berzosertib (Selleckchem), bestatin methyl ester (Abcam), blasticidin (InvivoGen), bortezomib (Selleckchem), cycloheximide (Sigma-Aldrich), doramapimod (Cayman), doxycycline (Thermo Fisher Scientific), doxorubicin (Selleckchem), etoposide (Calbiochem), ISRIB (Selleckchem), JNK-IN-8 (Selleckchem), KU-60019 (Selleckchem), Lactimidomycin (Sigma-Aldrich), MG-132 (Selleckchem), MLN4924 (MedChem Express), MLN7243 (ChemieTek), PF-3644022 (Sigma-Aldrich), SB202190 (Sigma-Aldrich), and Vx-765 (Selleckchem).

### Nanobody library generation

To raise heavy chain-only antibodies against NLRP1^PYD^, two alpacas were five times immunized with 200 µg His-NLRP1^PYD^ using Imject^™^ Alum Adjuvant (Thermo Fisher Scientific) according to locally authorized protocols. VHH plasmid libraries in the M13 phagemid vector pD (pJSC) were generated as described before (Schmidt et al., 2016b). In brief, RNA from peripheral blood lymphocytes was extracted and used as a template to generate cDNA using three sets of primers (random hexamers, oligo(dT), and primers specific for the constant region of the alpaca heavy chain gene). VHH coding sequences were amplified by PCR using VHH-specific primers, cut with AscI and NotI, and ligated into pJSC linearized with the same restriction enzymes. *E. coli* TG1 cells (Agilent) were electroporated with the ligation reactions and the obtained ampicillin-resistant colonies were harvested, pooled, and stored as glycerol stocks.

### Nanobody identification

NLRP1^PYD^ VHHs were obtained by phage display and panning with a protocol modified from Schmidt et al. (Schmidt et al., 2016b). *E. coli* TG1 cells containing the VHH library were infected with helper phage VCSM13 to produce phages displaying the encoded VHHs as pIII fusion proteins. Phages in the supernatant were purified and concentrated by precipitation. Phages presenting NLRP1^PYD^-specific VHHs were enriched using GST-NLRP1^PYD^ immobilized to magnetic Pierce glutathione beads (Thermo Fisher Scientific). The retained phages were used to infect *E. coli* ER2738 and subjected to a second round of panning. 96 *E. coli* ER2837 colonies yielded in the second panning were grown in 96-well plates and VHH expression was induced with IPTG. VHHs leaked into the supernatant were tested for specificity using ELISA plates coated with control protein GST or GST-NRLP1^PYD^. Bound VHHs were detected with HRP-coupled rabbit anti-E-Tag antibodies (Bethyl), and the chromogenic substrate TMB. Reactions were stopped with 1 M HCl and absorption at 450 nm was recorded. Positive candidates were sequenced and representative nanobodies were cloned into bacterial expression vectors for further analysis.

### ELISA

To test nanobody candidates, GST-NLRP1^PYD^ or GST in PBS were immobilized on ELISA plates at a concentration of 1μg/mL overnight. Serial 10-fold dilutions of HA-tagged nanobodies in 10% FBS/PBS were incubated with the immobilized antigen, followed by incubation with HRP-coupled anti-HA (clone 6E2, 1:5000, Cell Signaling), and the chromogenic substrate TMB. Reactions were stopped with 1 M HCl and absorption was measured at 450 nm.

### Flow cytometry-based quantification of inflammasome assembly

To quantify the assembly of ASC-EGFP specks or recruitment of C1C-EGFP to ASC specks (‘C1C specks’), cells were typically treated in 24-wells and analyzed by flow cytometry. We seeded 2.5·10^5^ HEK^NLRP1+ASC^ (or other derivatives of HEK 293T), or 10^5^ N/TERT-1^C1C-EGFP^ (or other derivatives of N/TERT-1) per well and cultivated cells overnight in the absence of antibiotics. Cells were stimulated for 6 or 20 h in DMEM (10% FBS) (HEK 293T) or Keratinocyte SFM (BPE, EGF, CaCl_2_, 100 µM Vx-765) (N/TERT-1), harvested by trypsinization, fixed in 4% formaldehyde, and analyzed using BD FACSCanto and BD LSRFortessa SORP flow cytometers, recording area, width, and height of the EGFP signal of single cells. Where indicated, cells were pre-treated with small compound inhibitors for 30 min before stimulation. UV stimulation was performed by irradiating cells in tissue culture plates (without lid) in a Bio-Link UVB irradiation system equipped with 5x 8 Watt T-8.M tubes emitting UVB at 312 nm for 3 min, followed by cultivation for 20 h. For infection experiments, cells were infected for 1 h in serum-free medium at the indicated multiplicity of infection (MOI), ranging from 1-50. Medium was subsequently replaced with full medium (with indicated inhibitors), and cells cultivated for another 19 h. To transiently overexpress proteins in HEK-based reporter cells, cells were transfected using Lipofectamine 2000 (Thermo Fisher Scientific) or PEI Max (Polysciences), medium replaced after 4 h, and cells harvested 20 h post transfection. Dox-inducible expression in lentivirus-generated cell lines was initiated by cultivating cells in 1 µg/mL dox for 6 or 20 h. Where indicated, fixed and permeabilized cells were stained with anti-HA (1:1000), anti-FLAG (1:300), anti-phospho-p38 (1:500), anti-gamma H2A.X (phospho S139) (1:500), anti-dsRNA (1:500), or anti-VSV G (1:1,000) in Intracellular Staining Permeabilization Wash Buffer (Biolegend) combined with Alexa Fluor (AF) 405, or AF647-coupled, highly cross-absorbed secondary antibodies (Thermo Fisher Scientific).

### Cytokine quantification by HTRF

To quantify IL-1β secretion, N/TERT-1-derived cells were stimulated as described for flow cytometry experiments in the absence and presence of Vx-765. IL-1β was quantified using the Human IL1 beta HTRF kit (Ciscbio) according to the manufacturer’s instructions. Supernatants for the quantification of IL-1β levels after inducible expression of VHL-VHH fusions were collected from 5·10^4^ cells per well in 96-well plates.

### Cell death quantification by LDH release

Cell death by pyroptosis was quantified after inducible expression of VHL-VHH fusions in N/TERT-1 cells. Supernatants for the quantification of lactate dehydrogenase activity were collected from 5·10^4^ cells per well in 96-well plates, including control samples in which cells were lysed in 1% Triton X-100. Lactate dehydrogenase activity was quantified using the LDH Cytotoxicity Detection Kit (TaKaRa) according to the manufacturer’s instructions. Cell death was normalized to the values of Triton X-100-lysed cells after removal of medium background.

### Microscopy

To generate microscopy samples, cells were seeded on 12 mm cover slips in 24-well plates and otherwise stimulated as described for flow cytometry experiments. Cells were fixed in 4% formaldehyde and stained for DNA using Hoechst 33342 (1:5,000). Images were recorded with a Zeiss Observer.Z1 wide field microscope, or a Leica SP8 Lightning confocal microscope. For live cell imaging experiments, N/TERT-1^C1C-EGFP^ cells were seeded in tissue culture-treated CellCarrier-96 Ultra Microplates (Perkin Elmer) and treated with the indicated stimuli in the presence of 3.3 µg/mL propidium iodide (PI). Cells were were cultivated at 37°C, 5% CO2 and images recorded every 20 min for 20 hours using a Zeiss Observer Z1 wide-field microscope with 20X PlanApochromat objective (NA = 0.8).

### Immunoblot

To generate immunoblot samples, 10^6^ cells were lysed in 200 uL 1x SDS-PAGE samples buffer. Proteins were separated by SDS-PAGE, transferred to PVDF membrane, and probed with the indicated antibodies, followed by HRP-coupled secondary antibodies (Thermo Fisher Scientific). Chemiluminescent signal of Western Lightning Plus-ECL (Perkin Elmer) was detected using a Fusion Advancer imaging system (Vilber).

### In vitro kinase assay

For *in vitro* kinase assays, 0.2 µM kinase was incubated with 7 µM substrate and 0.2 mM ATP containing 0.45 mCi [32P]-γ-ATP/mL (Perkin Elmer) in kinase buffer (50 mM HEPES pH 7.6, 34 mM KCl, 7 mM MgCl2, 2.5 mM dithiothreitol, 5 mM β-glycerol phosphate). Reactions were incubated for 30 min at 30° C and 300 rpm, and stopped by addition of EDTA to a final concentration of 50 mM. Samples were spotted onto Amersham Protran nitrocellulose membrane (GE Healthcare), followed by three washing steps for 5 min each with PBS. Counts per minute were determined in a Beckman Liquid Scintillation Counter (Beckman-Coulter) for 1 min. Measurements were performed in quadruplicate and represented as mean with standard deviation (SD).

## Acknowledgments

We thank Janett Wieseler for excellent technical support, and Patrick Schumachers for initial virus preparation and infection experiments. We would like to acknowledge the assistance of Andreas Dolf and the Flow Cytometry Core Facility at the Institute of Experimental Immunology, Medical Faculty at the University of Bonn. We are grateful to Gabor Horvath and the Microscopy Core Facility of the Medical Faculty at the University of Bonn for providing help and services, supported by the Deutsche Forschungsgemainschaft (DFG, German Research Foundation) for projects 13123509 (SFB 670) and 266686698. We would like to thank Paul-Albert Koenig and the Core Faciltiy Nanobodies at the Medical Faculty of the University of Bonn for providing the infrastructure for camelid immunizations. We are grateful to Jonathan L. Schmid-Burgk for help with the validation of knockout cell lines.

The presented work was supported by the following funding agencies:

Deutsche Forschungsgemeinschaft (DFG, German Research Foundation) grant GRK2168-272482170 (FIS)

Deutsche Forschungsgemeinschaft (DFG, German Research Foundation) grant SFB1403-414786233 (EL and FIS)

Deutsche Forschungsgemeinschaft (DFG, German Research Foundation) grant SFB1454-432325352 (EL),

Deutsche Forschungsgemeinschaft (DFG, German Research Foundation) grant TRR237-369799452 (BMK, EL, and FIS)

Deutsche Forschungsgemeinschaft (DFG, German Research Foundation) grant SPP1923-429513120 (EL and FIS)

Deutsche Forschungsgemeinschaft (DFG, German Research Foundation) Emmy Noether Programme 322568668 (FIS)

Deutsche Forschungsgemeinschaft (DFG, German Research Foundation) grant GE 976/9-2 (MG)

Deutsche Forschungsgemeinschaft (DFG, German Research Foundation) Germany’s Excellence Strategy – EXC2151–390873048 (EL, FIS, and MG)

Klaus Tschira Boost Fund (KT07) (FIS)

## Declaration interests

EL and FIS are cofounders and shareholders of Dioscure Therapeutics SE and Odyssey Therapeutics; MG and EL are a cofounder and shareholder of IFM Therapeutics; FIS is a consultant and shareholder to IFM Therapeutics. The other authors declare no competing interest.

