## Supplemental Information for "P38 kinases mediate NLRP1 inflammasome activation after ribotoxic stress response and virus infection"

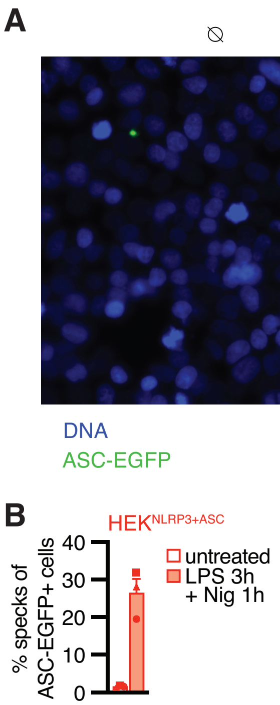


**Fig. S1. Reconstituted reporter cell lines recapitulate NLRP1 inflammasome assembly. (A)** HEK$^{NLRP1+ASC}$ cells were seeded on cover slips and stimulated with 30 µM talabostat for 20 h. Fixed cells were stained for DNA and representative images recorded by wide-field fluorescence microscopy. Scale bars represent 50 µm. (**B**) HEK$^{NLRP3+ASC}$ cells were treated with 200 ng/mL LPS for 3 h and 10 µM nigericin (Nig) for 1 h, followed by quantification of ASC specks by flow cytometry as described in Fig. 1A. Data represents average values (with individual data points) from three independent experiments ± SEM.

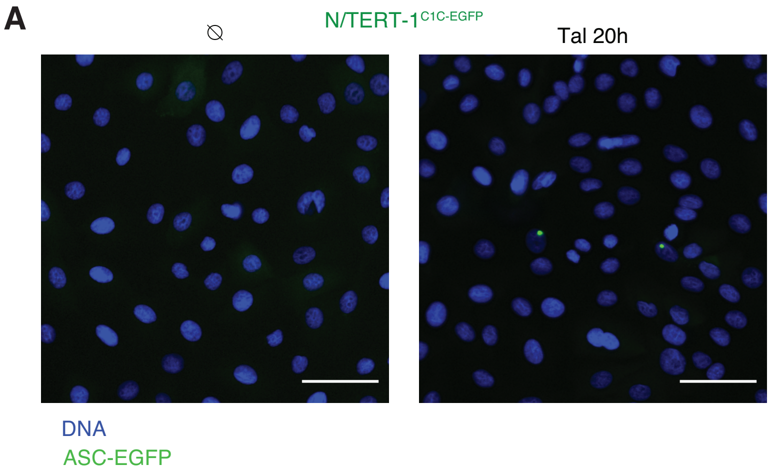


**Fig. S2. Reporter keratinocyte cell lines recapitulate NLRP1 inflammasome assembly.**

(**A**) N/TERT-1$^{C1C\text{-}EGFP}$ cells were seeded on cover slips and stimulated with 30 µM talabostat for 20 h. Fixed cells were stained for DNA and representative images recorded by wide-field fluorescence microscopy. Scale bars represent 50 µm.

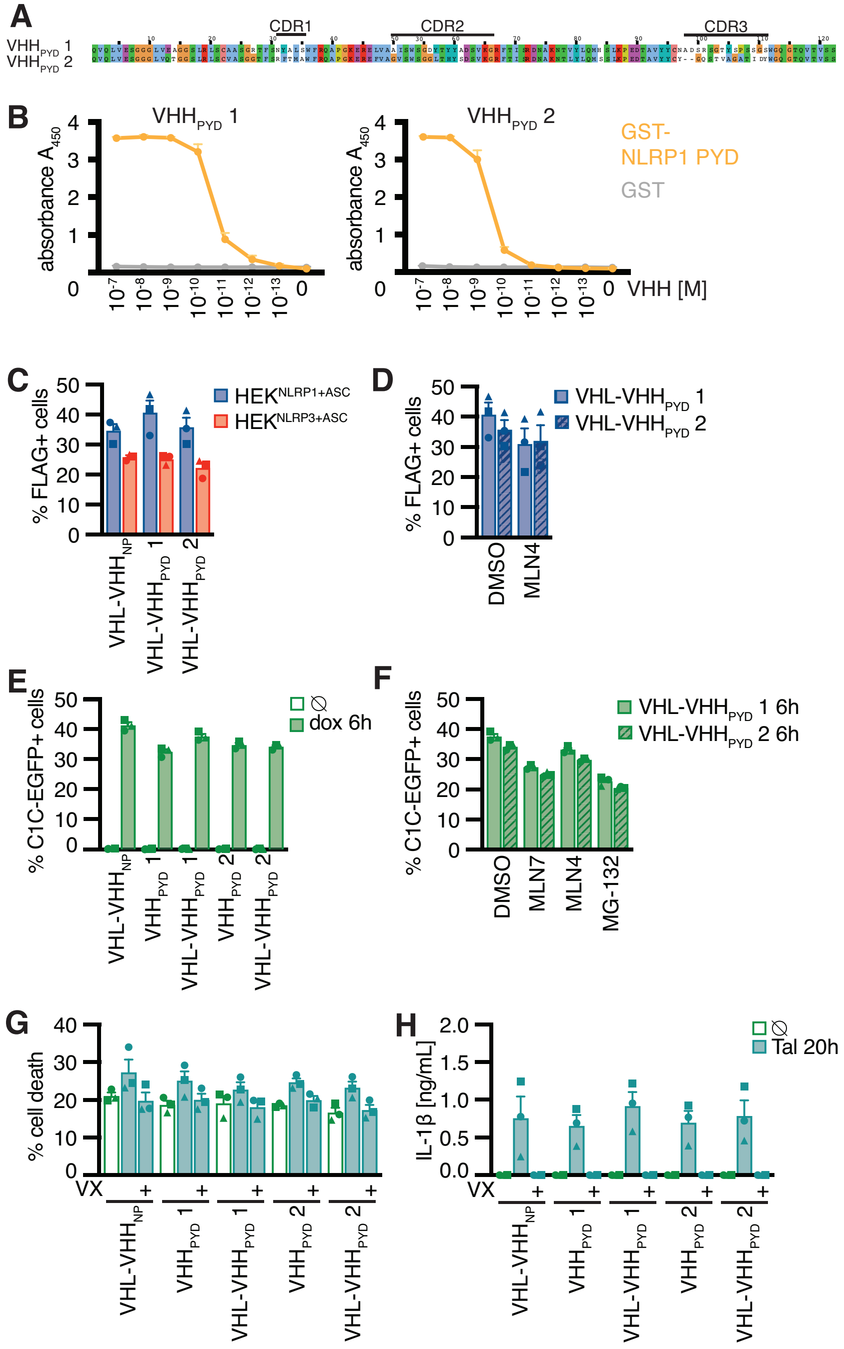


**Fig. S3. Ubiquitination of NLRP1$^{PYD}$ is sufficient for activation of human NLRP1.**
(**A**) Alignment of NLRP1$^{PYD}$ VHHs with indicated complementarity determining regions (CDRs). **(B)** Binding of indicated concentration of HA-tagged VHHs to immobilized GST-NLRP1$^{PYD}$ or control protein GST was quantified by ELISA. (**C, D**) HEK$^{NLRP1+ASC}$ and HEK$^{NLRP3+ASC}$ from the experiments described in Fig. 3B (C) and Fig. 3C (D) were stained for FLAG and the fraction of VHL-VHH-FLAG expressing cells quantified by flow cytometry. (**E, F**) C1C-EGFP expression of N/TERT-1 cells from the experiments described in Fig. 3D (E) and Fig. 3E (F) was quantified by flow cytometry. (**G, H**) N/TERT-1 keratinocytes encoding C1C-EGFP and the indicated VHH or VHL-VHH constructs were stimulated with 30 μM talabostat in the absence of dox. Cell death was evaluated by LDH release (G), and IL-1β secretion by HTRF (H). Experiments were

performed in parallel to experiments shown in Fig. 3. D, G, and H. Data represents average values (with individual data points) from three independent experiments ± SEM.

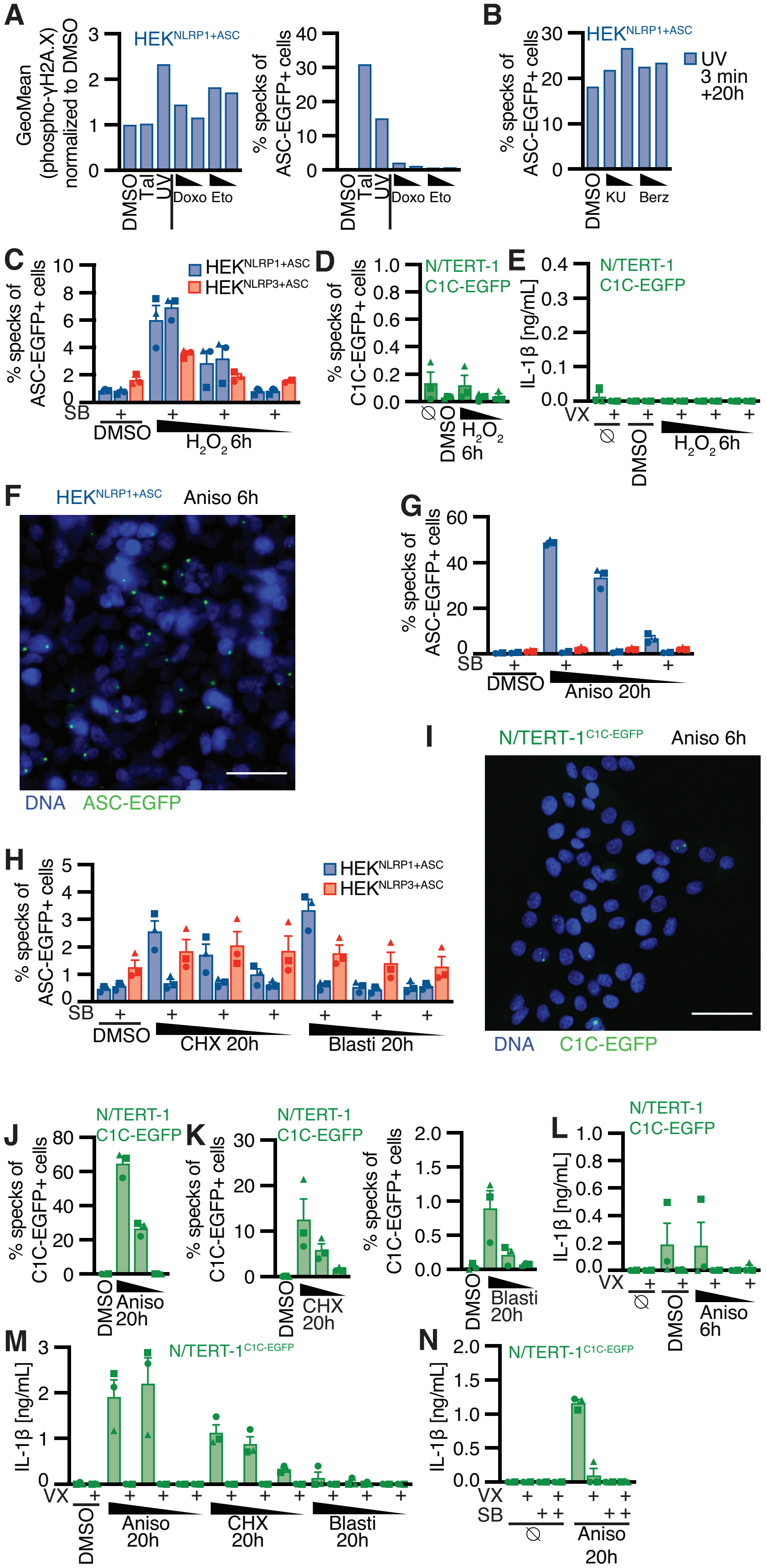
A
B
C
D
E
F
G
H
I
J
K
L
M
N
HEK NLRP1+ASC
HEK NLRP3+ASC
N/TERT-1 C1C-EGFP
GeoMean (phospho-γH2A.X) normalized to DMSO
% specks of ASC-EGFP+ cells
% specks of C1C-EGFP+ cells
IL-1β [ng/mL]
UV 3 min +20h
DMSO
Tal
UV
Doxo
Eto
KU
Berz
SB
VX
H2O2 6h
Aniso 6h
Aniso 20h
CHX 20h
Blasti 20h
DNA
ASC-EGFP
C1C-EGFP

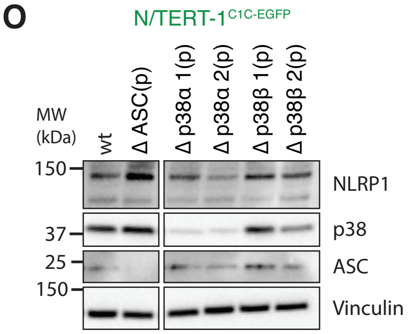


**Fig. S4. Human NLRP1 is activated by the ribotoxic stress response.** (**A**) HEK$^{NLRP1+ASC}$ cells were stimulated with 30 µM talabostat, UV for 3 min, 20 or 2 µM doxorubicin (Doxo), or 100 and 25 µM etoposide (Eto) and harvested after 20 h. Cells were stained for phospho-γH2A.X and analyzed for DNA damage markers (left) or ASC-EGFP specking (right) by flow cytometry. (**B**) HEK$^{NLRP1+ASC}$ cells were stimulated with UV for 3 min and subsequently cultivated in the presence of 10 or 1 µM KU-60019 (KU), or 1 and 0.1 µM Berzosertib (Berz) for 20 h, followed by quantification of ASC specks as in (A). (**C-E**) HEK$^{NLRP1+ASC}$, HEK$^{NLRP3+ASC}$ (C), or N/TERT-1$^{C1C-EGFP}$ (D-E) cells were treated with 1.5, 0.3, and 0.06 mM $H_2O_2$ for 6 h. Cells were stimulated in the presence of 20 µM SB where indicated; N/TERT-1$^{C1C-EGFP}$ cells were stimulated in the presence of 100 µM Vx-765 (VX) in (D), and where indicated in (E). Specks and secreted IL-1β were quantified by flow cytometry and HTRF, respectively. **(F, I)** HEK$^{NLRP1+ASC}$ (F) or N/TERT-1$^{C1C-EGFP}$ (I) cells were seeded on cover slips, treated with 15 µM anisomycin (Aniso) for 6 h, fixed, and stained for DNA. Representative images were recorded by wide-field fluorescence microscopy. Scale bars represent 50 µm. (**G, J**) HEK$^{NLRP1+ASC}$, HEK$^{NLRP3+ASC}$ (G) or N/TERT-1$^{C1C-EGFP}$ (J) cells were treated with 15, 1.5, and 0.15 µM anisomycin for 20 h; stimulation was performed in the presence of 20 µM SB where indicated in G, and in the presence of 100 µM VX in (J). (**H, K**) HEK$^{NLRP1+ASC}$, HEK$^{NLRP3+ASC}$ (H) or N/TERT-1$^{C1C-EGFP}$ (K) cells were treated with 1000, 200, and 40 µM cycloheximide (CHX), or 40, 4, and 0.8 µg/mL blasticidin S (Blasti); stimulation was performed in the presence of 20 µM SB where indicated in (H), and in the presence of 100 µM VX in (K).

(**L-N**) N/TERT-1$^{C1C-EGFP}$ cells were treated with anisomycin, CHX, and blasticidin dilutions as in (J) and (K), where indicated in the presence of 100 µM VX and/or 20 µM SB. IL-1β in the supernatant was quantified by HTRF. (**O**) Immunoblot analysis of cell lysates of wt N/TERT-1$^{C1C-EGFP}$ and the indicated polyclonal knockouts. Data represents average values (with individual data points) from three independent experiments ± SEM. Microscopy images are representative of three independent experiments.

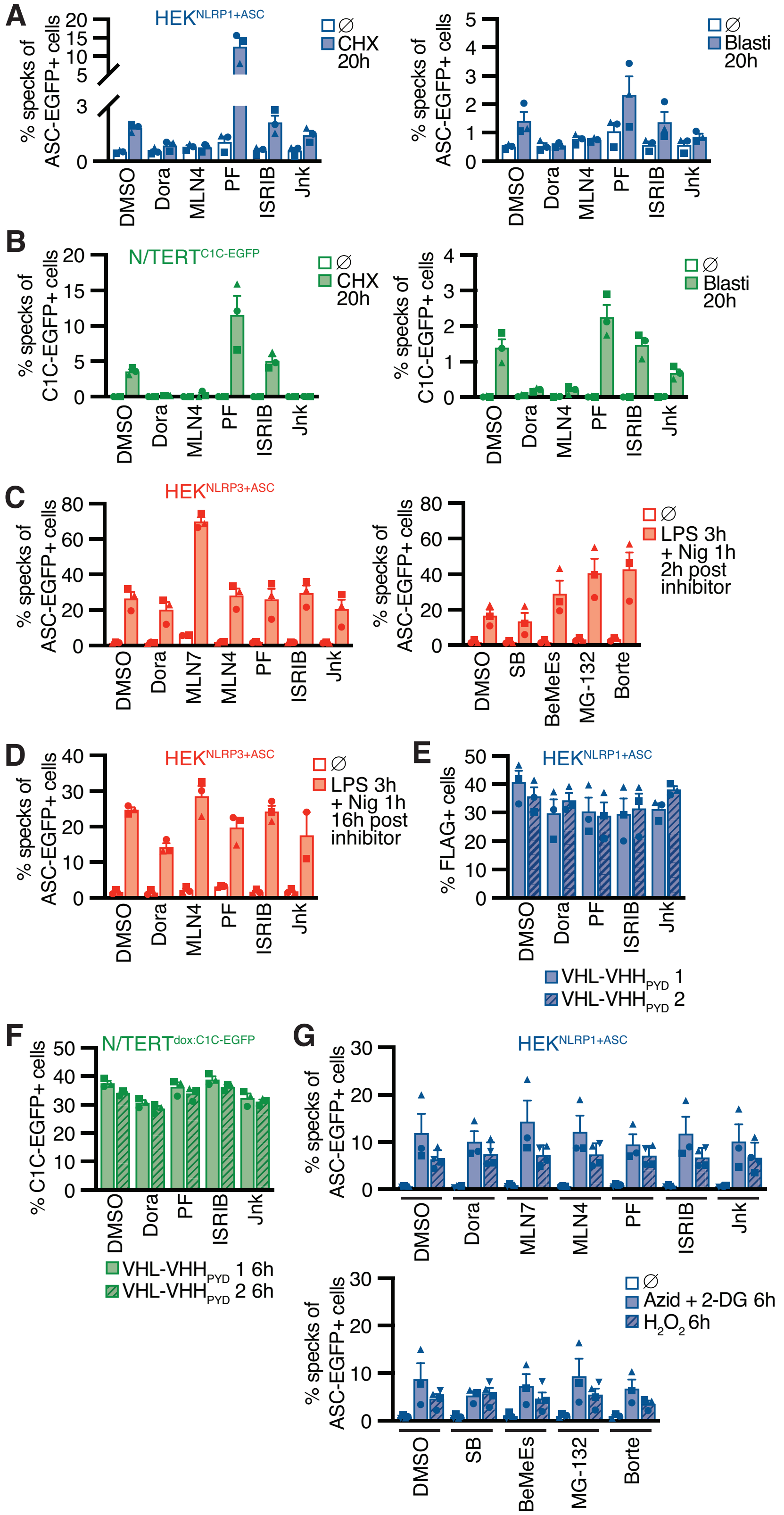


**Fig. S5. NLRP1 activation by ribotoxic stress response relies on the ubiquitination machinery and proteasomes.** HEK$^{NLRP1+ASC}$ (A), N/TERT-1$^{C1C-EGFP}$ (B), HEK$^{NLRP3+ASC}$ (C, D) were stimulated with 1 mM CHX, 20 µg/mL blasticidin (A, B), or 200 ng/mL LPS for 3 h and 10 µM Nig for 1 h (C, D) in the presence of the indicated inhibitors as in Fig. 5. Note that LPS+Nigericin stimulation was performed towards the end of inhibitor treatment to evaluate NLRP3 responses after 6 h (C) and 20 h (D) in the presence of the indicated inhibitors. Experiments were done in parallel to data shown in Fig. 5, A and C. (**E**) VHL-VHH-FLAG expression of transiently transfected $HEK^{NLRP1+ASC}$ cells from Figure 5D was quantified by flow cytometry. (**F**) Inducible C1C-EGFP expression of N/TERT-1 keratinocytes from Figure 5G was quantified by flow cytometry. (**G**) $HEK^{NLRP1+ASC}$ were stimulated with 10 mM azide and 50 mM 2-DG or 1.5 mM $H_2O_2$ in the presence of the indicated inhibitors as described in Fig. 5. Data represents average values (with individual data points) from three independent experiments ± SEM.

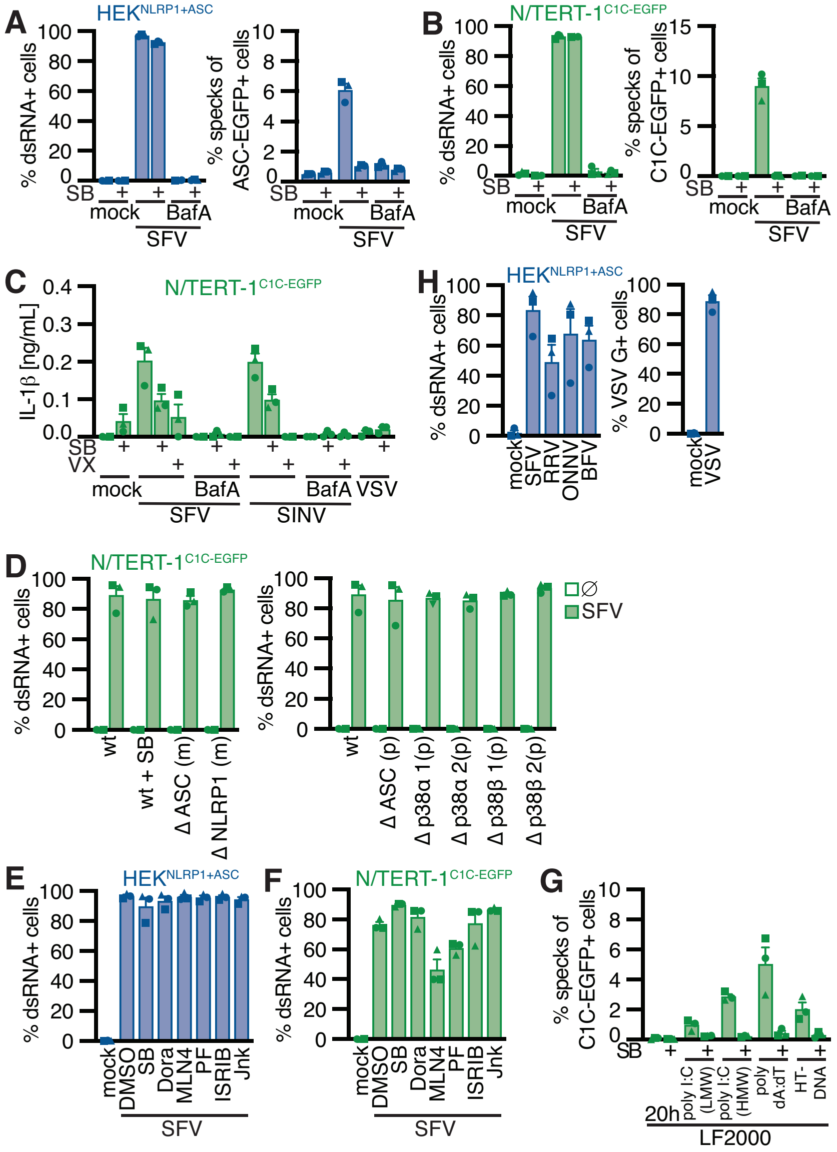


**Fig. S6. Alphavirus infection activates NLRP1 in a p38-dependent manner.** (**A, B**) HEK$^{\text{NLRP1+ASC}}$ (A) or N/TERT-1$^{\text{C1C-EGFP}}$ (B) treated with or without 100 nM bafilomycin A1 (BafA) and 20 µM SB where indicated (and 100 µM VX in B) were infected with Semiliki Forest virus (SFV) for 20 h. Cells were stained for dsRNA, followed by quantification of infection and ASC specks by flow cytometry. (**C**) N/TERT-1$^{\text{C1C-EGFP}}$ cells treated with the indicated drugs as in (A) were infected with SFV, SINV, or VSV for 20 h and IL-1β secretion quantified by HTRF. (**D-F**) N/TERT-1$^{\text{C1C-EGFP}}$ (D, F) and HEK$^{\text{NLRP1+ASC}}$ cells (E) from experiments in Fig. 6, C-E, were stained for dsRNA and infection quantified by flow cytometry. (**G**) N/TERT-1$^{\text{C1C-EGFP}}$ cells treated 100 µM VX, as well as 20 µM SB where indicated, were transfected with 1 µg/mL of the indicated nucleic acids using LF2000 and cultivated for 20 h; C1C-EGFP speck assembly was quantified by flow cytometry. Experiments in Fig. 2H and fig. S6G were done in parallel and values for stimulated cells in the absence of SB are displayed in both panels. Data represents average values (with individual data points) from three independent experiments ± SEM.

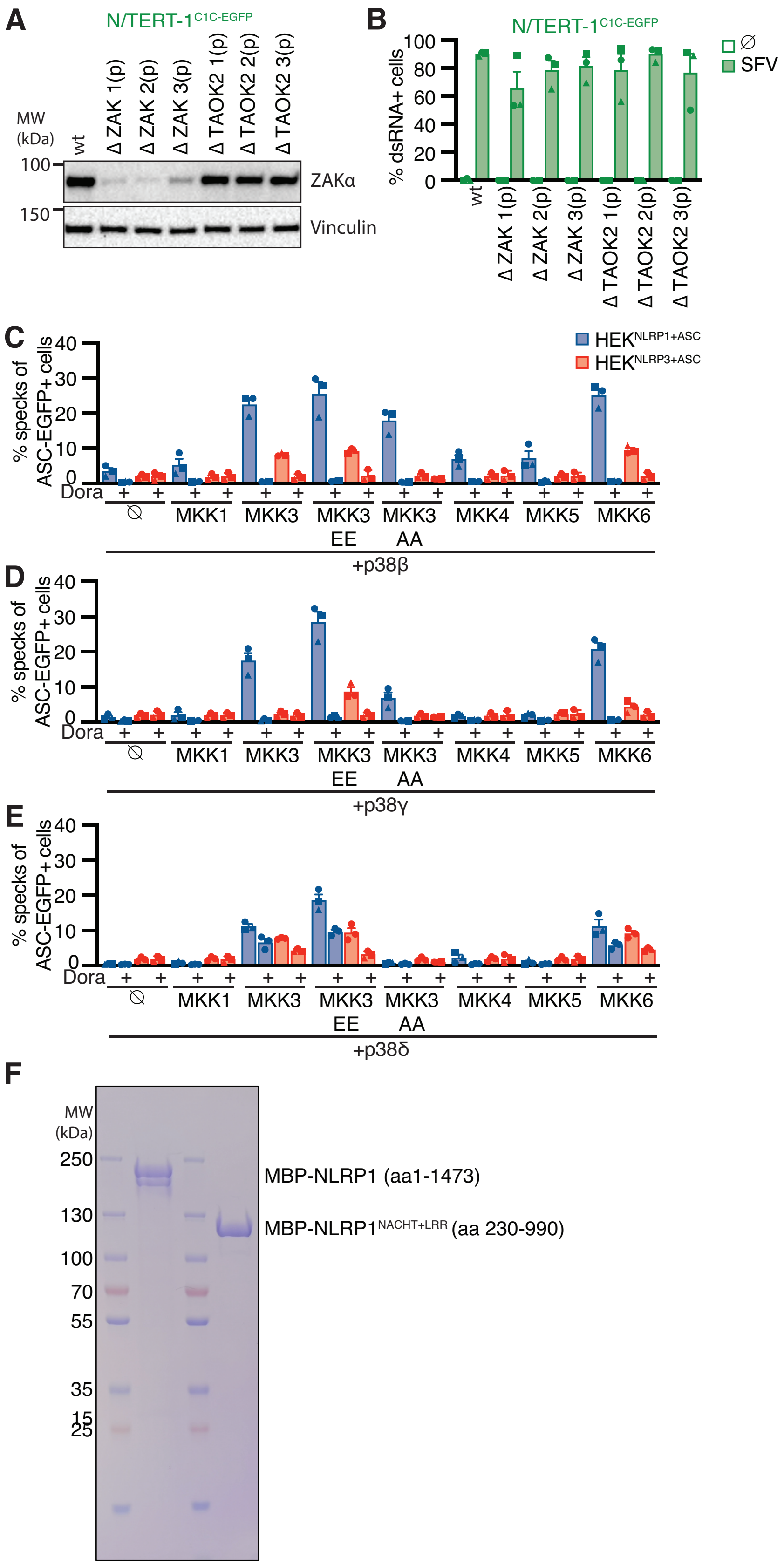
A
N/TERT-1 C1C-EGFP
MW (kDa)
wt
Δ ZAK 1(p)
Δ ZAK 2(p)
Δ ZAK 3(p)
Δ TAOK2 1(p)
Δ TAOK2 2(p)
Δ TAOK2 3(p)
100
150
ZAKα
Vinculin
B
N/TERT-1 C1C-EGFP
% dsRNA+ cells
∅
SFV
C
% specks of ASC-EGFP+ cells
HEK NLRP1+ASC
HEK NLRP3+ASC
Dora
MKK1
MKK3
MKK3 EE
MKK3 AA
MKK4
MKK5
MKK6
+p38β
D
+p38γ
E
+p38δ
F
MW (kDa)
250
130
100
70
55
35
15
25
MBP-NLRP1 (aa1-1473)
MBP-NLRP1 NACHT+LRR (aa 230-990)

**Fig. S7. P38 activation is sufficient for NLRP1 activation.** (**A**) Immunoblot analysis of cell lysates of wt N/TERT-1$^{\text{C1C-EGFP}}$ and the indicated polyclonal knockouts with the indicated antibodies. (**B**) N/TERT-1$^{\text{C1C-EGFP}}$ cells from experiments in Fig. 7A, were stained for dsRNA and infection quantified by flow cytometry. (**C-E**) HEK$^{\text{NLRP1+ASC}}$ or HEK$^{\text{NLRP3+ASC}}$ cells were transiently transfected with the indicated expression vectors and analyzed as described for Fig. 7, B-D. (**F**) SDS-PAGE gel of purified MBP-NLRP1 and MBP-NLRP1$^{\text{NACHT+LRR}}$ used as kinase substrate in Fig. 7E.

**Movie S1. Human NLRP1 is activated by the ribotoxic stress response.**

N/TERT-1$^{C1C\text{-}EGFP}$ cells in the presence of propidium iodide (PI) were treated with DMSO and recorded by wide-field microscopy over time.

**Movie S2. Human NLRP1 is activated by the ribotoxic stress response.**

N/TERT-1$^{C1C\text{-}EGFP}$ cells in the presence of propidium iodide (PI) were treated with 30 µM talabostat and recorded by wide-field microscopy over time.

**Movie S3. Human NLRP1 is activated by the ribotoxic stress response.**

N/TERT-1$^{C1C\text{-}EGFP}$ cells in the presence of propidium iodide (PI) were treated with 15 µM anisomycin and recorded by wide-field microscopy over time.

**Movie S4. Human NLRP1 is activated by the ribotoxic stress response.**

N/TERT-1$^{C1C\text{-}EGFP}$ cells in the presence of propidium iodide (PI) were treated with 2 µM Lacti and recorded by wide-field microscopy over time.
